# In-vivo whole-cortex marker of excitation-inhibition ratio indexes cortical maturation and cognitive ability in youth

**DOI:** 10.1101/2023.06.22.546023

**Authors:** Shaoshi Zhang, Bart Larsen, Valerie J. Sydnor, Tianchu Zeng, Lijun An, Xiaoxuan Yan, Ru Kong, Xiaolu Kong, Ruben C. Gur, Raquel E. Gur, Tyler M. Moore, Daniel H. Wolf, Avram J Holmes, Yapei Xie, Juan Helen Zhou, Marielle V Fortier, Ai Peng Tan, Peter Gluckman, Yap Seng Chong, Michael J Meaney, Gustavo Deco, Theodore D. Satterthwaite, B.T. Thomas Yeo

## Abstract

A balanced excitation-inhibition ratio (E/I ratio) is critical for healthy brain function. Normative development of cortex-wide E/I ratio remains unknown. Here we non-invasively estimate a putative marker of whole-cortex E/I ratio by fitting a large-scale biophysically-plausible circuit model to resting-state functional MRI (fMRI) data. We first confirm that our model generates realistic brain dynamics in the Human Connectome Project. Next, we show that the estimated E/I ratio marker is sensitive to the GABA-agonist benzodiazepine alprazolam during fMRI. Alprazolam-induced E/I changes are spatially consistent with positron emission tomography measurement of benzodiazepine receptor density. We then investigate the relationship between the E/I ratio marker and neurodevelopment. We find that the E/I ratio marker declines heterogeneously across the cerebral cortex during youth, with the greatest reduction occurring in sensorimotor systems relative to association systems. Importantly, among children with the same chronological age, a lower E/I ratio marker (especially in association cortex) is linked to better cognitive performance. This result is replicated across North American (8.2 to 23.0 years old) and Asian (7.2 to 7.9 years old) cohorts, suggesting that a more mature E/I ratio indexes improved cognition during normative development. Overall, our findings open the door to studying how disrupted E/I trajectories may lead to cognitive dysfunction in psychopathology that emerges during youth.

**Significance:** Healthy brain function requires a delicate balance of neural excitation (E) and inhibition (I). In animals, this balance – the E/I ratio – is known to decrease with the maturation of inhibitory circuitry during healthy development. However, in humans, the normative development of cortex-wide E/I ratio remains unclear. Here, we use a biophysical model and non-invasive brain scans to estimate a marker of E/I ratio. Spatial changes in our E/I ratio marker are consistent with a drug that decreases E/I ratio. We also find that our cortex-wide E/I ratio marker decreases during development. Furthermore, North American and Asian children with lower E/I ratio, especially in higher-order cortex, have better cognitive performance. Overall, E/I ratio is a potential index of healthy neurocognitive development.

## 1. Introduction

Healthy brain function requires a delicate balance between neural excitation (E) and inhibition (I) (1–4). This balance – the E/I ratio – is refined during critical developmental periods of heightened experience-dependent plasticity (5,6). E/I imbalances during critical developmental periods are thought to contribute to the etiology of many psychiatric disorders (7,8) and confer vulnerability to cognitive deficits (9,10). Here we capitalize on advances in biophysically plausible large-scale circuit models to chart the normative development of cortex-wide E/I ratio and uncover links to cognition.

Human cortical development unfolds hierarchically – sensory systems mature earlier, while association systems follow a more protracted developmental course extending through adolescence (11,12). A potential mechanism driving this hierarchical development might be the temporal progression of critical plasticity periods along the sensorimotor-to-association axis (13–16). More specifically, the maturation of GABAergic inhibitory circuitry involving parvalbumin positive (PV) interneurons suppresses stimulus-irrelevant activity, yielding a higher signal-to-noise ratio (13). The maturation of PV interneurons also modulates long-term potentiation by enforcing a narrower spike integration window (17). Overall, the maturation of the inhibitory circuitry facilitates the experience-dependent pruning of excitatory pyramidal neuronal connections via the Hebbian mechanism, triggering a critical plasticity period (18–20). Therefore, a hallmark feature of the critical period development is a reduction in the E/I ratio (21,22). While the hierarchical progression of inhibitory development is documented in animal models (14,23), it is unclear if the same mechanisms exist in humans, extend to the evolutionarily expanded association cortex and impact cognitive ability.

Studying E/I ratio development *in-vivo* in humans is challenging due to limitations in non-invasive neuroimaging techniques. MR spectroscopy studies suggest changes in the balance of excitatory and inhibitory neurotransmitter levels in single brain regions during development (24,25). A recent study used a machine learning marker trained with pharmacological-fMRI data to provide evidence of E/I ratio reduction in the association cortex during development (26). However, these past studies were limited to partial portions of the cortex, so normative development of cortex-wide E/I ratio remains unclear. Indirect estimates of whole-cortex E/I ratio have provided insights into autism spectrum disorder in adults and Alzheimer’s Disease (27–29), but these approaches mostly lack a direct mapping to an underlying biophysically-plausible model of excitatory and inhibitory dynamics.

Biophysically-plausible large-scale circuit models of coupled brain regions have provided mechanistic insights into spontaneous brain dynamics (30–32). However, most large-scale circuit models assume that local synaptic properties are spatially uniform across brain regions (9,27,33), which lacks biological plausibility. Indeed, spatial heterogeneity in excitatory and inhibitory cell types (34–36) might be a driver of large-scale brain dynamics (37,38). Studies have shown that incorporating spatial heterogeneity across local synaptic parameters generates more realistic spontaneous brain dynamics (39,40). Our previous study (41) demonstrated that parameterizing local synaptic parameters with anatomical and functional gradients led to dramatically more realistic brain dynamics in adults. However, we utilized a large-scale circuit model (42) that did not differentiate among excitatory and inhibitory neural populations, so the E/I ratio could not be derived.

Here we investigate the development of cortical E/I ratio over youth and its association with cognitive ability. We apply our previous approach (41) to the feedback inhibition control (FIC) model with coupled excitatory and inhibitory neuronal populations (33). The resulting parameteric FIC (pFIC) model is used to derive a potential marker of E/I ratio. We first confirm that the pFIC model yields realistic brain dynamics in healthy young adults from the Human Connectome Project (HCP; 43). Using a pharmacological fMRI dataset (44), we show that the E/I ratio marker is sensitive to E/I ratio reduction induced by the GABA-agonist alprazolam. Then, using the Philadelphia Neurodevelopmental Cohort (PNC; 45,46), we find that the E/I ratio declines across the cortex during youth. Furthermore, a lower E/I ratio indexes greater cognitive ability, with the strongest relationships observed in association cortex. We generalize the link between E/I ratio and cognitive ability in a younger GUSTO (Growing Up in Singapore with Healthy Outcomes) cohort (47). Overall, our study suggests that E/I ratio maturation might be a driver of healthy neurocognitive development during youth.

## 2. Results

### 2.1 Overview

We first evaluated the optimization of the spatially heterogeneous parametric feedback inhibition control (pFIC) model in the HCP dataset (**Figure 1A**). The biological plausibility of the estimated marker of excitation-inhibition (E/I) ratio was then evaluated using pharmacological fMRI involving GABAergic benzodiazepine alprazolam. Finally, we investigated developmental changes and cognitive effects of E/I ratio in the PNC dataset. Assocations with cognition were replicated in the GUSTO cohort.

**Figure 1.**
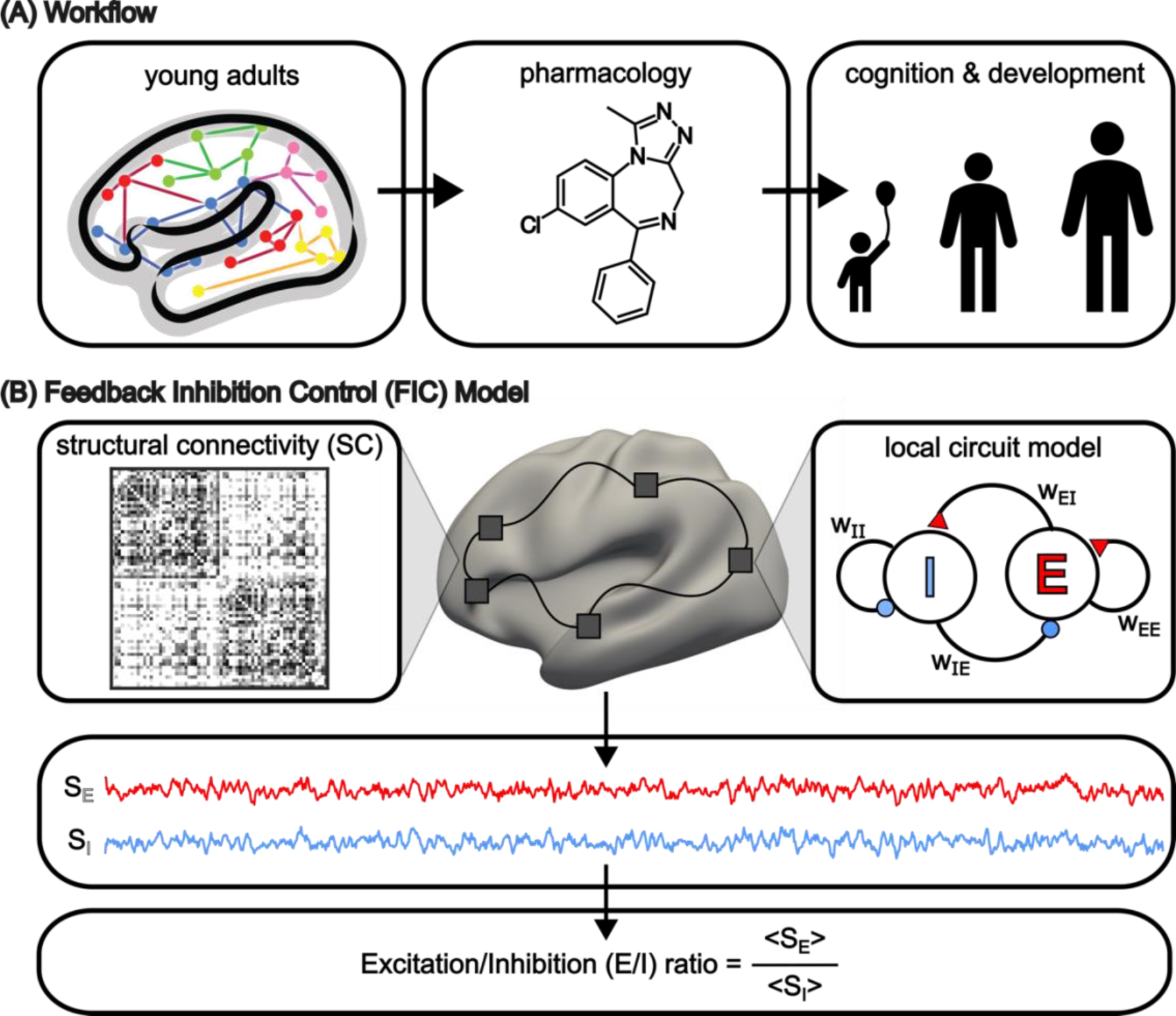
Workflow and schematic of the parametric feedback inhibition control (pFIC) model. (A) Young adults from the Human Connectome Project (HCP) were used to evaluate the optimization of the spatially heterogeneous pFIC model. Pharmacological fMRI with benzodiazepine alprazolam was then used to evaluate the biological plausibility of the estimated E/I ratio. Next, the pFIC model was used to investigate the development of cortex-wide E/I ratio and its association with cognitive ability in the PNC dataset. Cognitive associations were replicated in a sample of seven-year-olds from the GUSTO cohort. HCP logo is used with permission from the HCP team. (B) The FIC model (33) is a neural mass model obtained by mean field reduction of spiking neuronal network models. The FIC model consists of differential equations at each cortical region governing the neural dynamics of excitatory and inhibitory neuronal populations (“E” and “I” respectively in the right panel). A red triangle indicates an excitatory connection. A blue circle indicates an inhibitory connection. “w_xy_” indicates the connection strength from neuronal population x to neuronal population y. For example, “w_IE_” indicates the connection strength from the inhibitory population to the excitatory population. The regional models are connected by excitatory connections parameterized by a structural connectivity (SC) matrix. For a given set of model parameters, time courses of excitatory (S_E_) and inhibitory (S_I_) synaptic gating variables (representing the fraction of open channels) can be simulated. The excitation-inhibition ratio (E/I ratio) was defined as the ratio between the temporal average of S_E_ and S_I_. Local synaptic parameters were estimated using the same approach as our previous study (41). We refer to the resulting model as the pFIC model.

### 2.2 Optimization of the parametric feedback inhibition control (pFIC) model

We randomly divided 1004 HCP participants (43,48) into 3 non-overlapping training (N = 335), validation (N = 335), and test (N = 334) sets. The Desikan–Killiany anatomical parcellation (49) with 68 cortical regions of interest (ROIs) was used to generate group-level structural connectivity (SC), static functional connectivity (FC), and functional connectivity dynamics (FCD) from the training, validation, and test sets separately. To compute FCD for each fMRI run, a 68 × 68 FC matrix was computed for each sliding window of length ∼60 seconds. The 68 × 68 FC matrices were then correlated across the 1118 windows, yielding a 1118 × 1118 FCD matrix (41). The FCD matrix has been shown to reflect temporal fluctuations in resting-state FC that are not captured by static FC (50,51). See Supplementary Methods S2 for details.

The FIC model (33) is a neural mass model obtained by mean field reduction of a spiking neuronal network model (52,53). The model comprises ordinary differential equations (ODEs) at each cortical region describing the dynamics of excitatory and inhibitory neuronal populations (**Figure 1B** top panel). The local dynamics are driven by recurrent connections within separate excitatory and inhibitory populations, as well as connections between excitatory and inhibitory populations. Greater excitatory-to-excitatory recurrent strength (w_EE_) and smaller inhibitory-to-excitatory connection strength (w_IE_) amplify synaptic currents of the excitatory population. Similarly, greater excitatory-to-inhibitory connection strength (w_EI_) and smaller inhibitory-to-inhibitory recurrent strength (w_II_) amplify synaptic currents of the inhibitory population. Neuronal noise in each cortical region is controlled by the noise amplitude *σ*. Finally, the excitatory populations of the regional local models are connected via the SC matrix, scaled by a global constant *G*.

Following previous studies (33,39), w_II_ was set to one and w_IE_ was automatically set to maintain a uniform baseline excitatory firing rate of around 3Hz. Excitatory-to-excitatory recurrent strength (w_EE_), excitatory-to-inhibitory connection strength (w_EI_), regional noise amplitude (*σ*) and the SC scaling constant (*G*) were estimated using our previous approach (41). More specifically, w_EE_, w_EI_ and *σ* were parameterized as a linear combination of the principal resting-state FC gradient (54) and T1w/T2w myelin estimate (55), resulting in 9 unknown linear coefficients and 1 unknown parameter *G*. We refer to the resulting model as the parametric FIC (pFIC) model.

The 10 pFIC parameters were estimated using the covariance matrix adaptation evolution strategy (CMA-ES) (56) by minimizing the difference between simulated and empirical fMRI data. More specifically, agreement between empirical and simulated FC matrices was defined as the Pearson’s correlation (*r*) between the *z*-transformed upper triangular entries of the two matrices. Larger *r* indicates more similar static FC. However, Pearson’s correlation does not account for scale difference, so we also computed the absolute difference (*d*) between the means of the empirical and simulated FC matrices (39). A smaller *d* indicates more similar static FC. The inclusion of *d* was necessary to prevent overly synchronized fMRI signals (**Figure S1**). Finally, we do not expect the brain states of two participants to be the same at any given timepoint *t* during the resting-state, i.e., there is no temporal correspondence between participants in the resting-state. Because FCD was computed based on sliding window FC, there was similarly no temporal correspondence between simulated and empirical FCD matrices. Therefore, disagreement between the simulated and empirical FCD matrices was defined as the Kolmogorov–Smirnov (*KS*) distance, following previous studies (41,50). The KS distance was defined as the maximum distance between the cumulative distribution functions obtained by collapsing the upper triangular entries of simulated and empirical FCD matrices, so no temporal correspondence was assumed (more details in Supplementary Methods S10). The overall cost was defined as (1 – *r*) + *d* + *KS*. A smaller cost indicates better agreement between simulated and empirical fMRI.

### 2.3 The pFIC model generates realistic fMRI dynamics

We first demonstrate that the parametrization of the local synaptic parameters with T1w/T2w and FC gradient led to more realistic brain dynamics than spatially homogeneous parameters **(Figure 2A)**. We applied CMA-ES to the HCP training set to generate 500 candidate model parameter sets. The 500 parameter sets were evaluated in the HCP validation set. The top 10 parameter sets from the HCP validation set were used to simulate FC and FCD using SC from the HCP test set, which were then compared with empirical FC and FCD from the HCP test set. A strong agreement between simulated and empirical FC (as well as between simulated and empirical FCD) would suggest that the pFIC model was able to generate realistic brain dynamics.

**Figure 2.**
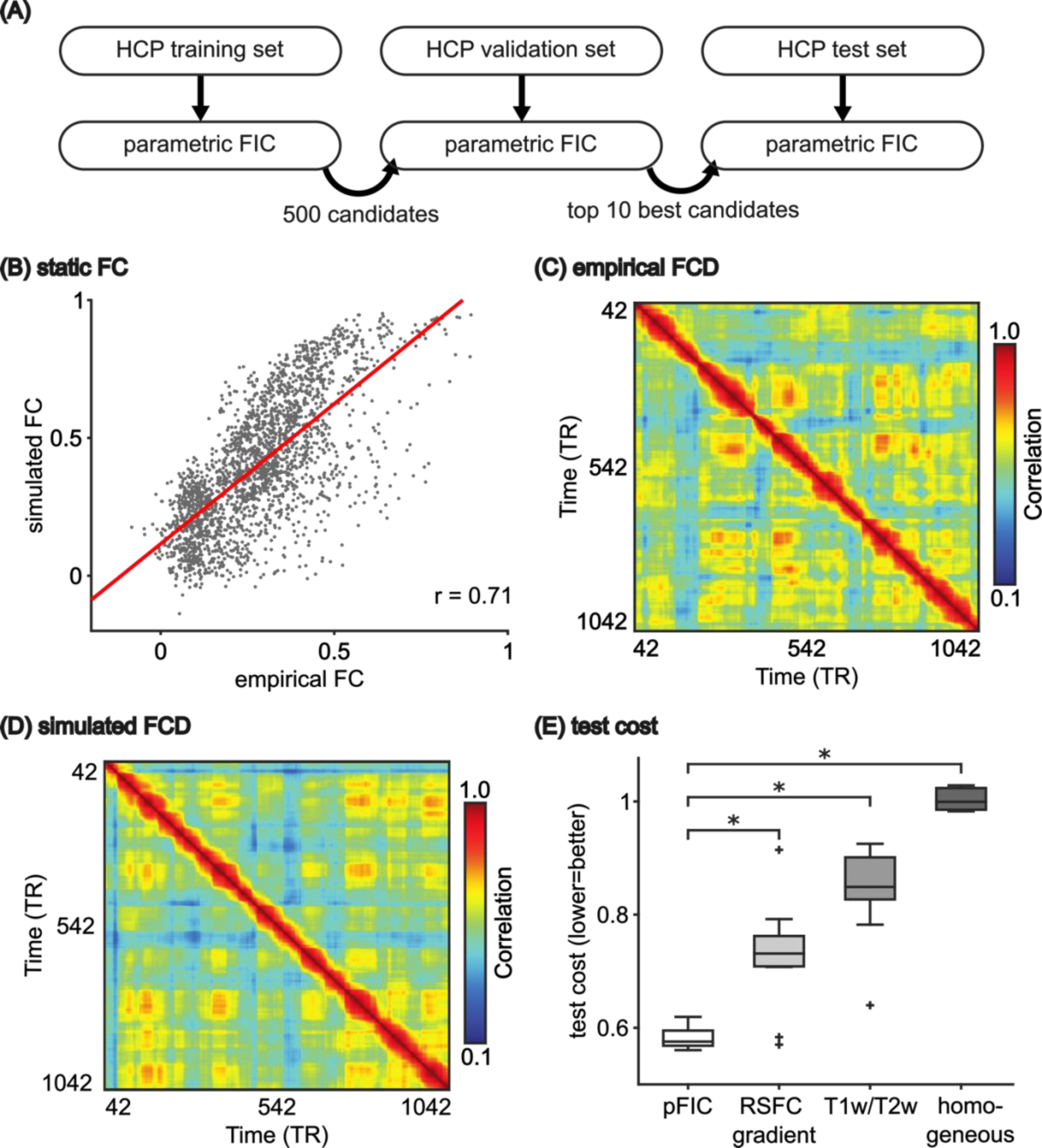
The parametric feedback inhibition control (pFIC) model generates more realistic fMRI dynamics than the spatially homogeneous FIC model. (A) The CMA-ES algorithm (41,56) was applied to the HCP training set to generate 500 sets of model parameters. The top 10 parameter sets from the validation set were evaluated in the test set. (B) Agreement (Pearson’s correlation) between empirical and simulated static FC in the HCP test set. (C) Empirical FCD from an HCP test participant. (D) Simulated FCD from the pFIC model using the best model parameters (from the validation set) and SC from the test set. (E) Total test cost of the pFIC model compared with three control conditions: (1) local synaptic parameters parameterized by only principal resting-state FC gradient, (2) local synaptic parameters parametrized by only T1w/T2w ratio map and (3) local synaptic parameters constrained to be spatially uniform. The boxes show the inter-quartile range (IQR) and the median. Whiskers indicate 1.5 IQR. Black crosses represent outliers. * indicates that the pFIC model achieved statistically better (lower) test cost.

**Figure 2B** visualizes the correlation between empirical and simulated FC in the HCP test set (based on the best model parameters from the validation set). Across the 10 best parameter sets from the validation set, correlation between empirical and simulated static FC was 0.71 ± 0.005 (mean ± std) in the test set. As a reference, correlation between empirical FC and SC in the test set was 0.48. On the other hand, the absolute difference between the means of the empirical and simulated FC matrices was 0.11 ± 0.015 in the test set. This suggests that the pFIC model was able to generate realistic FC.

**Figure 2C** shows the empirical FCD from a single run of a representative HCP test participant. **Figure 2D** shows the simulated FCD using the best model parameters (from the validation set) and SC from the test set. Off-diagonal red blocks in both empirical and simulated FCD indicated recurring FC patterns that were not simply due to temporal autocorrelation. Similarity in the amount of off-diagonal red blocks between empirical and simulated FCD suggests that the pFIC model was able to generate realistic FCD. Across the 10 best candidate sets from the validation set, the Kolmogorov-Smirnov (*KS*) distance between empirical and simulated FCD was 0.18 ± 0.028 in the HCP test set. Disagreement between simulated and empirical fMRI appeared more pronounced in posterior regions, but the pattern of disagreement was not correlated with the RSFC gradient or the T1w/T2w ratio map (**Figures S2 and S3**).

Overall, the pFIC model was able to generate realistic FC and FCD, yielding an overall cost of 0.58 ± 0.018 in the HCP test set (**Figure 2E**). Parameterizing model parameters with only the principal FC gradient or only T1w/T2w ratio map led to worse (higher) cost in the HCP test set (**Figure 2E**). Most large-scale circuit model studies assume spatially homogeneous parameters. When local synaptic parameters (w_EE_, w_EI_ and *σ*) were constrained to be uniform across brain regions (33,57) and optimized by CMA-ES, the cost was poor in the test set (**Figure 2E**). These results emphasize the importance of parameterizing local synaptic parameters with spatial gradients that smoothly varied from sensory-motor to association cortex. Consistent with our previous study (41), the T1w/T2w and FC gradient were complementary in the sense that combining the two spatial maps led to more realistic fMRI dynamics.

### 2.4 Estimated E/I ratio is sensitive to the effect of benzodiazepine alprazolam

In the previous section, we showed that the pFIC model could be effectively optimized to generate realistic fMRI dynamics. Here, we evaluated the biological plausibility of the estimated E/I ratio in a pharmacological-fMRI dataset (44) comprising 45 participants, who completed a placebo-controlled double-blind fMRI study with benzodiazepine alprazolam. Alprazolam is a benzodiazepine that enhances GABAergic signaling at GABA_A_ receptor sub-units, including *α*_1,2,3,5_ and *γ*_1−3_ (58,59). Alprazolam enhances GABAergic inhibitory signaling via positive allosteric modulation, thus reducing E/I ratio (60). Therefore, we hypothesized that the E/I ratio estimated with the pFIC model would be lower during the alprazolam condition compared with the placebo condition.

The 45 participants were equally divided into training, validation and test sets. Group-level SC, first principal FC gradient, and T1w/T2w ratio maps from the HCP dataset were used in the following analysis. For each experimental condition (placebo or alprazolam), 250 candidate parameter sets were generated from the condition’s training set. The top 10 parameter sets from the validation set were evaluated in the test set. The costs of the 10 parameter sets generalized well to the test set (**Figure S4**), suggesting that there was no overfitting in the validation set.

One challenge in analyzing this dataset was that the fMRI data was acquired with a limited field of view (FOV). Therefore, 26 out of 68 Desikan-Killiany ROIs with less than 50% coverage (**Figure S5**) were not considered during the estimation of the model parameters. The estimated model parameters were extrapolated to the entire cortex (see Supplementary Methods S11 for details) and used to simulate the excitatory (S_E_) and inhibitory (S_I_) time courses (**Figure 1B**). Motivated by rodent studies, the E/I ratio was defined as the ratio between the temporal average of S_E_ and S_I_ (61).

An E/I ratio contrast was computed by subtracting the E/I ratio estimated during the drug (alprazolam) session from the E/I ratio estimated during the placebo session. Since alprazolam is expected to reduce E/I ratio, we hypothesized the E/I ratio would be lower during the alprazolam condition, yielding a positive E/I ratio contrast. Consistent with our hypothesis, the E/I ratio contrasts of all regions were positive (**Figure 3B**). 67 out of 68 regions exhibited E/I ratio contrasts statistically different from zero after correcting for multiple comparisons with a false discovery rate (FDR) of *q* < 0.05. We note that there was no motion difference between the drug and placebo fMRI sessions (*p* > 0.1). These results suggest that the E/I ratio estimated by the pFIC model was sensitive to the pharmacological enhancement of inhibitory activities.

**Figure 3.**
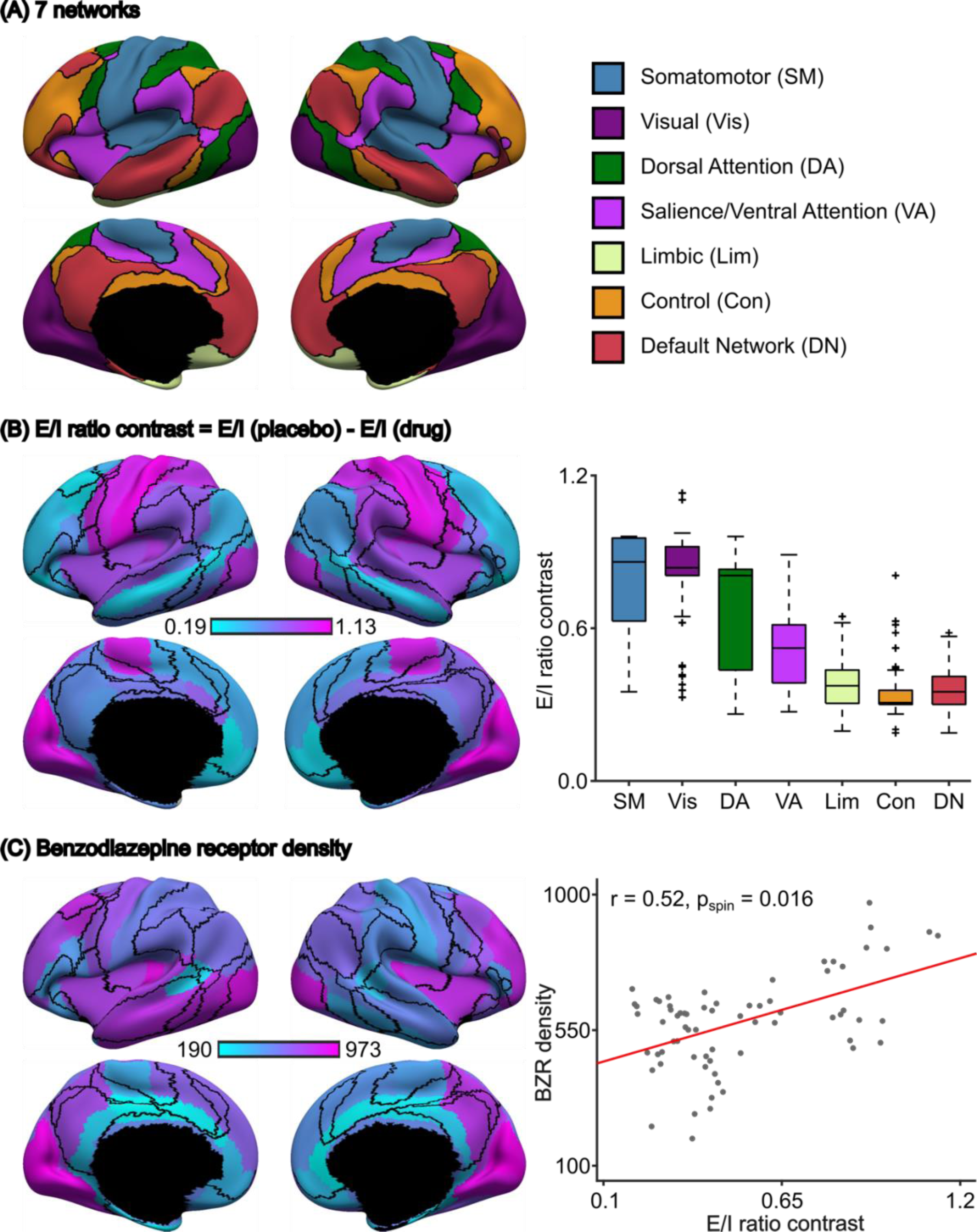
E/I ratio estimate is sensitive to the effect of benzodiazepine alprazolam. (A) Seven resting-state networks (64). (B) Left: Regional E/I ratio contrast overlaid with the boundaries (black) of seven resting-state networks. 67 out of 68 regions showed significant E/I ratio difference between placebo and drug sessions after FDR correction (*q* < 0.05). E/I ratio difference was greater than zero for all regions, consistent with lower E/I ratio during the alprazolam session. Right: E/I ratio differences exhibited a spatial gradient with higher differences in sensory-motor regions compared with regions in the control and default networks. The boxes show the inter-quartile range (IQR) and the median. Whiskers indicate 1.5 IQR. Black crosses represent outliers. (C) Spatial distribution of BZR density (pmol/ml) from *in-vivo* positron emission tomography in a separate group of participants (62). (D) Higher regional BZR density was associated with larger E/I ratio changes during the drug session (*r* = 0.52, two-tail spin test *p* = 0.016).

Since the distribution of benzodiazepine receptors (BZR) density is not spatially uniform (62), we hypothesized that the E/I ratio contrast would also not be spatially uniform, and would align with BZR density. Supporting this, we found that the E/I ratio contrast exhibited a spatial gradient with strongest effects in sensory-motor networks and weakest effects in control and default networks (**Figure 3B). Figure 3C** shows the spatial distribution of benzodiazepine receptors (BZR) density estimated from *in-vivo* positron emission tomography in a separate group of participants (62). Regions with greater BZR density exhibited greater reduction in E/I ratio during the drug session (*r* = 0.52; two-tail spin test *p* = 0.016; **Figure 3D**). Therefore, the spatial distribution of E/I ratio contrast was biologically plausible.

To evaluate robustness, we repeated the above analyses 5 times with different random splits of the 45 participants into training, validation and test sets. The results were similar across the 5 splits (**Figures S6 and S7**). Results from the most representative split were shown in **Figure 3**. Using this most representative split, we performed several additional sensitivity analyses. In the previous analyses, the acceptable excitatory firing rate was constrained to be between 2.7Hz and 3.3Hz. Relaxing the thresholds to between 2.5Hz and 3.5Hz yielded similar results (**Figure S8**). Changing the ROI coverage threshold from 50% to 60% also yielded similar results (**Figure S9**). We repeated the analysis using a 100-region homotopic functional parcellation (63), which also yielded similar results (**Figure S10**). Pairwise comparisons between the control analyses are found in **Figure S11**. Similar results were obtained with log-transformation or square root of BZR density (**Figure S12**).

### 2.5 The E/I ratio declines with development in youth

Having demonstrated that the E/I ratio estimates were sensitive to the alprazolam-induced enhancement of inhibitory activities, we next explored how the E/I ratio changes during development in the Philadelphia Neurodevelopmental Cohort (PNC; 45,46). We hypothesized that the estimated E/I ratio would decline with age.

After data preprocessing and quality control, we obtained a sample of 885 participants ages 8-23 years old (**Figure 4A**). Participants were sorted according to age and evenly divided into 29 age groups, so each group comprised 30 or 31 participants. Within each age group, 15 participants were randomly selected as the validation set, while the remaining participants were assigned to the training set. For each age group, 250 candidate model parameter sets were generated from the group’s training set using CMA-ES and evaluated in the group’s validation set; the parameter set with the lowest validation cost was used to estimate regional E/I ratio across the cortex.

**Figure 4.**
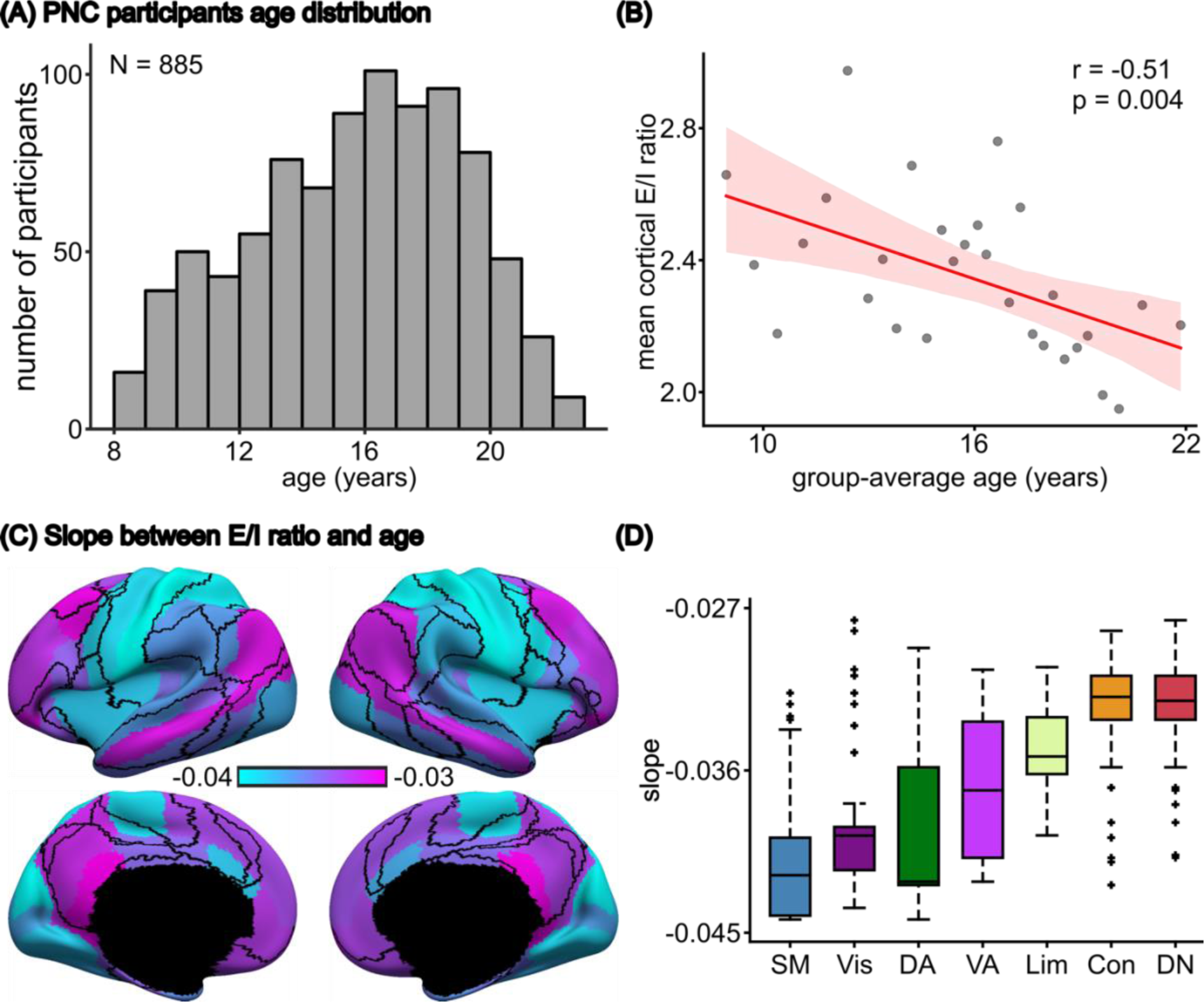
E/I ratio continuously declines throughout child and adolescent development. (A) Age distribution of 885 PNC participants (mean = 15.66, std = 3.36, min = 8.17, max = 23). (B) Participants in older age groups exhibited lower E/I ratio (*r* = −0.51, *p* = 0.004). Participants were divided into 29 non-overlapping age groups. There are 29 dots in the scatter plot, corresponding to the 29 age groups. The shaded area depicts 95% confidence interval of the linear relationship. (C) Spatial distribution of linear regression slope between regional E/I ratio and age. The values represent the rate of E/I ratio changes during development. All slopes were negative and significant (FDR *q* < 0.05). (D) The slopes exhibited a spatial gradient with sensory-motor networks showing the fastest E/I ratio reduction and association networks showing slower E/I ratio reduction. The boxes show the inter-quartile range (IQR) and the median. Whiskers indicate 1.5 IQR. Black crosses represent outliers.

We performed linear regression between age and mean cortical E/I ratio (i.e., E/I ratio averaged across the whole cortex), as well as between age and regional E/I ratio. Mean cortical E/I ratio declined throughout child and adolescent development (*r* = −0.51, *p* = 0.004; **Figure 4B**). This E/I ratio reduction was statistically significant for all cortical regions (FDR *q* < 0.05; **Figure 4C**). Furthermore, the rate of E/I ratio decrease exhibited a spatial gradient with sensory-motor regions exhibiting greater rate of E/I ratio decrease (i.e., more negative slope) compared with association networks (**Figure 4D**).

To evaluate the robustness of these effects, the PNC analyses were repeated 5 times with different splits of the participants (within each age group) into training and validation sets. The results were similar across the 5 random splits of the data (**Figures S13 and S14**). We conducted several additional sensitivity analyses using the most representitve split (which was shown in **Figure 4**). Relaxing the firing rate thresholds to between 2.5Hz and 3.5Hz yielded similar results (**Figure S15**), as did using a 100-region homotopic parcellation (63) (**Figure S16**). Pairwise comparisons between the control analyses are found in **Figure S17**. Finally, consistent with the literature, younger participants exhibited higher head motion during the fMRI scan. Therefore, as a control analysis, we regressed out mean framewise displacement from the E/I ratio estimates of each age group, yielding similar results (**Figure S18**).

### 2.6 Lower E/I ratio is associated with better cognition within the same age group

Having shown that older children exhibited lower E/I ratio (**Figure 3B**), we next evaluated the cognitive implications of such a decline in the E/I ratio as part of normative development. We hypothesize that a lower E/I ratio would be associated with better cognition. To test this hypothesis, we compared the E/I ratio of PNC participants who were matched on age but differed in cognitive performance.

Participants in the PNC completed the Penn Computerized Neurocognitive Battery (CNB), a 12-task battery that has been previously summarized using an overall (domain-general) measure of accuracy as well as three domain-specific factor scores (65). Participants were divided into 14 high-performance groups and 14 low-performance groups based on the overall accuracy measure. Each high-performance group was age-matched to a low-performance group (**Figure 5A**). Each low-performance or high-performance group comprised 31 or 32 participants. For each group, 15 participants were randomly assigned to the validation set, while the remaining participants were assigned to the training set. For each group, 250 candidate parameter sets were generated from the training set and the top parameter set from the validation set was used to estimate the E/I ratio; we compared the E/I ratio between the high and low performance groups.

**Figure 5.**
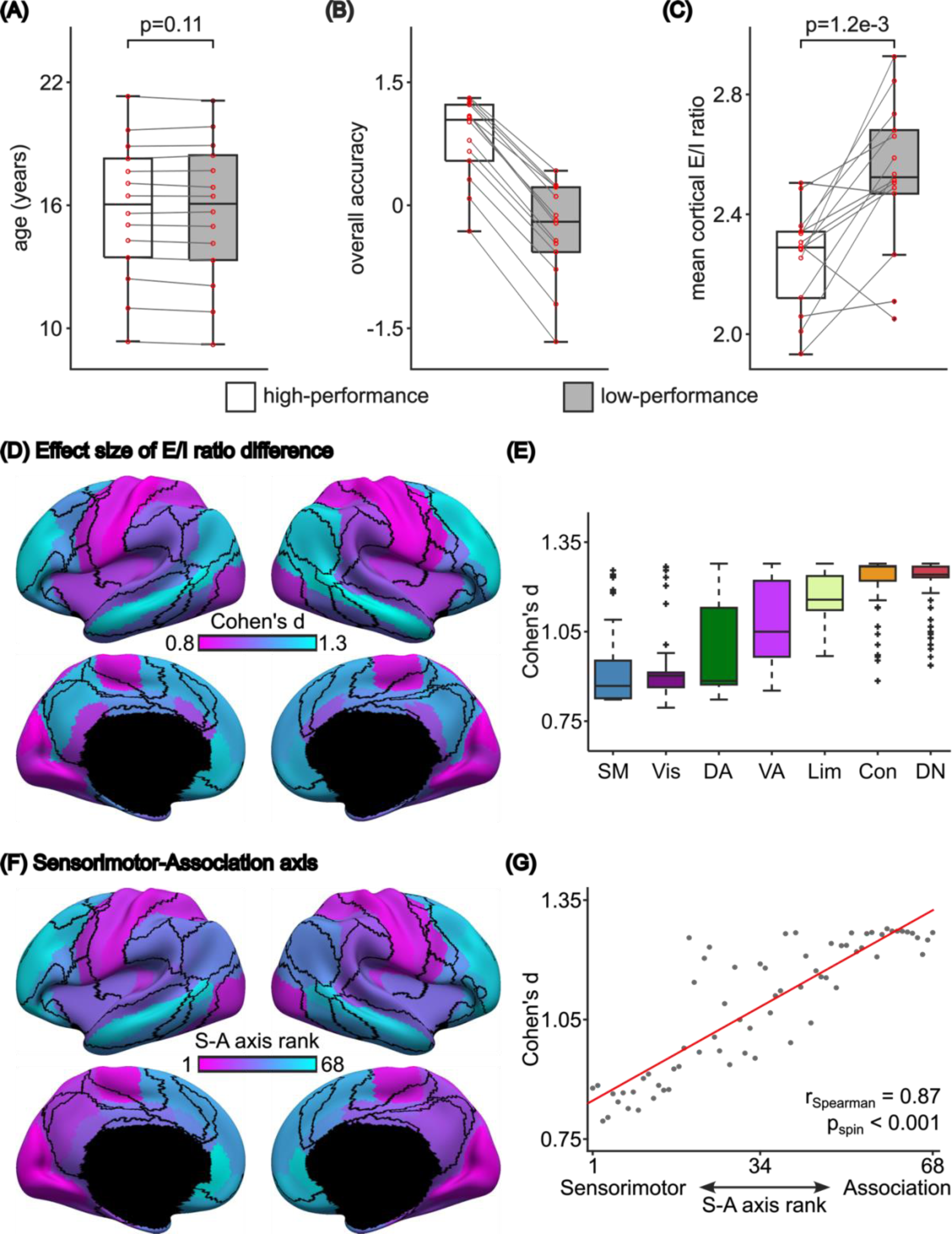
Lower E/I ratio is associated with better cognitive performance within the same age group in the PNC cohort. (A) Boxplots of age, (B) ‘overall accuracy’ and (C) mean cortical E/I ratio of high-performance and low-performance (overall accuracy) groups. The mean cortical E/I ratio of the high-performance group was significantly lower than that of the low-performance group (FDR *q* < 0.05). (D) Spatial distribution of effect size of regional E/I ratio difference between high-performance and low-performance groups. All regions were significant after FDR correction with *q* < 0.05. (E) Effect size of E/I ratio differences in cognition is larger in control and default networks compared with sensory-motor regions. The boxes show the inter-quartile range (IQR) and the median. Whiskers indicate 1.5 IQR. Black crosses represent outliers. (F) ROI rankings based on the sensorimotor-association (S-A) axis (66). Lower ranks were assigned to ROIs that were more towards the sensorimotor pole; higher ranks were assigned to ROIs that were more towards the association pole. (G) Agreement between effect size of E/I ratio difference and S-A axis rank. Spearman’s correlation *r* = 0.87, two-tailed spin test *p* < 0.001.

The high-performance group exhibited lower mean cortical E/I ratio than the low-performance group (two-tailed t-test *p* = 1.2 x 10^-3^; **Figure 5C**). There was no motion difference between high-performance and low-performance groups (*p* > 0.2). To test for domain specificity, we also compared the E/I ratio for the three domain-specifc factor scores (complex reasoning, memory and social cognition), but observed no statistical difference after correcting for multiple comparisons (**Figure S19**).

Having found global differences in the E/I ratio between the high and low cognitive performance groups, we next evaluated regional effects (**Figure 5D**). We found that E/I ratio differences between low-performance and high-performance groups were larger in control and default networks, compared with sensory-motor regions (**Figure 5E**; all FDR *q* <0.05). Notably, the effect sizes of these regional differences in the E/I ratio aligned well with the sensorimotor-association (S-A) axis of cortical organization (66), such that effect sizes were lowest at the sensorimotor pole and largest at the association pole (**Figure 5F**). Spearman’s correlation between effect sizes and S-A axis ranks was *r* = 0.87 (two-tailed spin test *p* < 0.001; **Figure 5G**). Overall, these results suggest that a more mature E/I ratio – especially in higher-order association cortex – is linked to more mature cognition.

To evaluate the robustness of these results, we repeated these analyses 5 times with different random training-validation splits of participants within each high-performance group and each low-performance group. The results were similar across the 5 splits (**Figures S20 to S24**; the most representative split is displayed in **Figure 5**). Within the most representative split, we found that relaxing the thresholds to between 2.5Hz and 3.5Hz yielded similar results (**Figure S25**) as did use of a 100-region homotopic functional parcellation (63) (**Figure S26**). Pairwise comparisons between the control analyses are found in **Figure S27**.

### 2.7 Results generalize to a younger Asian cohort

As a final step, we evaluated whether the link between the E/I ratio and cognition generalized to a group of younger participants of different anscestry. This was motivated by recent concerns that relationships between resting-fMRI and behavior may not generalize well across ethnic groups (67). We utilized the Growing Up in Singapore with Healthy Outcomes (GUSTO) dataset (47), which included 154 participants (after quality control) with a mean age of 7.5 years. An overall cognitive performance score was obtained by a principal component analysis of five cognitive tests. Participants were then divided into groups with high and low cognitive performance. The ages were well-matched between the high and low-performance groups (**Figures 6A and 6B**). There was no motion difference between the high and low-performance groups during the fMRI scans (*p* > 0.1).

**Figure 6.**
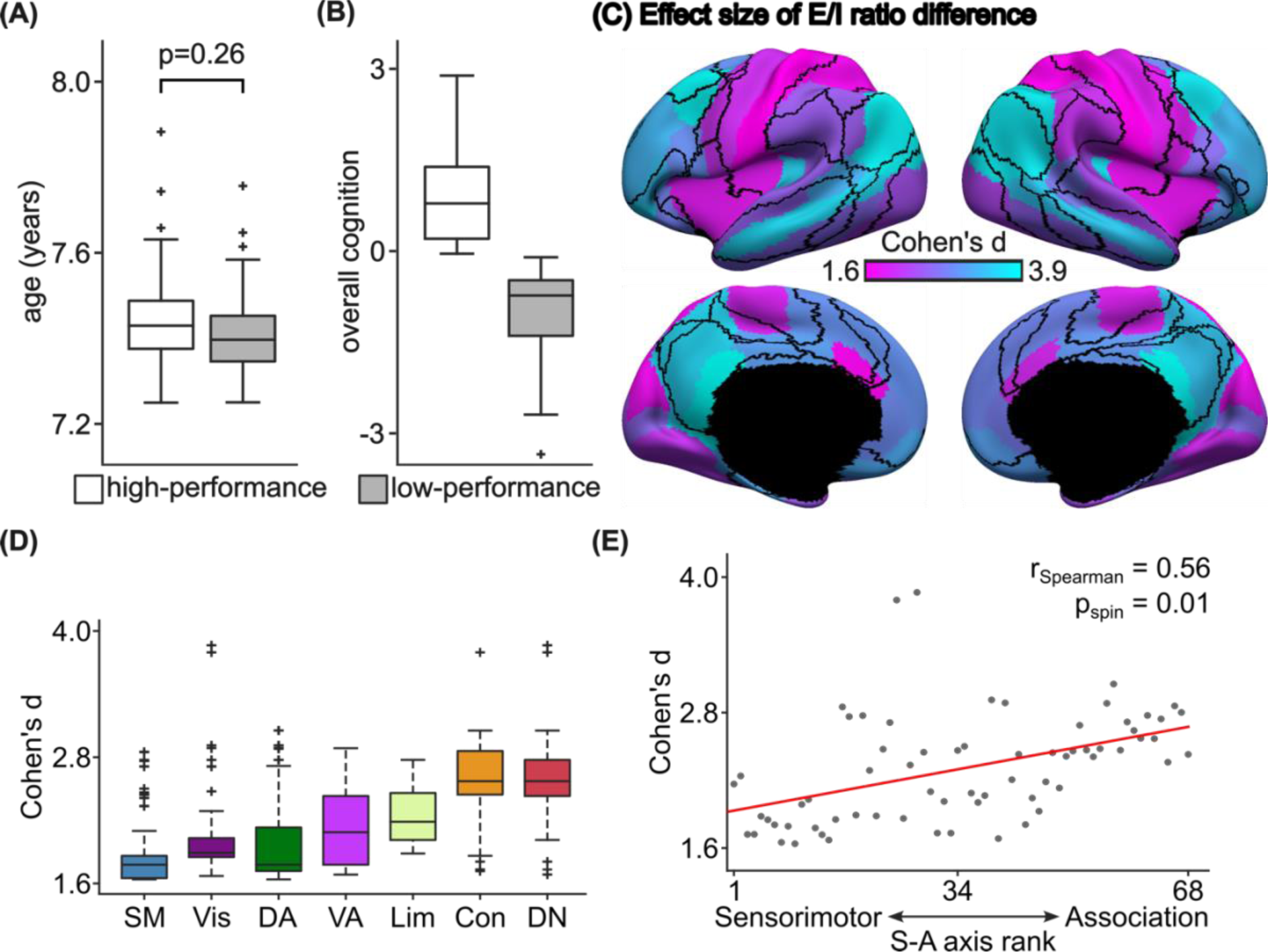
Lower E/I ratio is associated with better cognitive performance within the same age group in the GUSTO dataset. (A) Boxplots of age for high-performance and low-performance groups. (B) Overall cognition of high-performance and low-performance groups. (C) Spatial distribution of effect size of E/I ratio difference between low-performance group and high-performance group. (D) Effect size of E/I ratio differences is larger in control and default networks compared with sensory-motor regions. The boxes show the inter-quartile range (IQR) and the median. Whiskers indicate 1.5 IQR. Black crosses represent outliers. (E) Agreement between effect size of E/I ratio difference and S-A axis rank. Spearman’s correlation *r* = 0.56, two-tailed spin test *p* = 0.01.

Replicating PNC results, we found that the high-performance group exhibited a lower E/I ratio in higher-order association cortex than the low-performance group (**Figures 6C**). Differences were largest in the default and control networks (**Figure 6D**). Statistical significance was evaluated using a permutation test, where the null distribution was constructed by randomly assigning participants into high or low-performance groups, and then re-estimating the E/I ratio. We note that only 29 (out of 68) regions were significant after FDR correction with *q* < 0.05. These 29 regions were all in association cortex. By contrast, differences in the E/I ratio between cognitive performance groups was largely not significant in sensory-motor networks. As in the PNC, we found the effect sizes of these cognitive differences aligned with the S-A axis (*r* = 0.56, two-tailed spin test *p* = 0.01; **Figure 6E**). These results in a younger Asian cohort emphasize the robustness and generalizability of our findings.

## 3. Discussion

We first established that the pFIC model could generate realistic fMRI dynamics in a large adult dataset. We then demonstrated that our E/I ratio marker was sensitive to increased inhibitory activity induced by benzodiazepine alprazolam. In a large developmental sample from North America, we found that the E/I ratio marker decreased with age. We also demonstrated that a lower E/I ratio marker – reflective of more mature cortex – was associated with better cognitive performance, particularly in transmodal association cortex. Critically, these findings generalized to a younger Asian cohort. Together, our findings provide evidence that refinements in the cortical E/I ratio persist into adolescence, suggesting that prolonged E/I-linked developmental plasticity in the association cortex supports continued neurocognitive development. We speculate that insufficient refinement of the E/I ratio during development may create a vulnerability to cognitive deficits, with potentially important implications for transdiagnostic psychopathology.

The E/I ratio is challenging to be non-invasively investigated in humans. Post-mortem studies have established how the expressions of E/I relevant genes vary across the cortex (35,68,69). On the other hand, there is a lack of a direct mapping of *in-vivo* neuroimaging signals with excitatory and inhibitory neurobiology, as well as constrained spatial coverage and specificity of available E/I techniques (24–26). Here, we capitalized on recent developments in biologically interpretable computational modeling of cortical circuits to gain insight into the E/I ratio from fMRI data. We fitted a large-scale circuit model with interacting excitatory and inhibitory populations to resting-state fMRI and calculated the E/I ratio from the time courses of excitatory and inhibitory synaptic gating variables. Our E/I ratio marker captured reductions in the E/I ratio induced by alprazolam, a positive allosteric modulator that increases the effectiveness of GABAergic signaling (70). Furthermore, the spatial pattern of benzodiazepine-related E/I reductions described by the model was correlated with the distribution of benzodiazepine-sensitive GABA receptors from positron emission tomography (62). Interestingly, one of the pharmacological targets of benzodiazepines is GABA_A_ *α*_1_receptors (70). Increases in GABA signaling at GABA_A_ *α*_1_receptors have been shown to trigger the onset of developmental critical periods in animal models (71), indicating that the pFIC model is well equipped to study development-linked changes in inhibitory signaling in the human brain.

We found that the E/I ratio decreased across the cortex throughout child and adolescent development. E/I ratio declined with age across all cortical systems, but the magnitude of decline varied along a unimodal-transmodal cortical hierarchy. Specifically, by age 22, the E/I ratio had declined the most in unimodal sensory territories, such as the visual and somatomotor systems, and the least in transmodal systems like the default and frontoparietal control systems. Differences in E/I development across these systems may be linked to differences in their maturational time courses. The development of cortical inhibitory circuitry is well-established as a central mechanism that controls the timing and progression of critical periods of development (21,72). Initially, the development of inhibitory circuitry lags behind that of excitatory pyramidal cells, leading to an early increase in the local E/I ratio (73). Later, experience and evoked activity stimulate the development of inhibitory circuitry (21) — particularly fast-spiking parvalbumin positive interneurons and GABA_A_ *α*_1_ receptors — which begins to reduce the E/I ratio, facilitating experience-dependent plasticity and triggering the opening of the critical period window (22,71,72). As the critical period progresses, excitatory synapses are pruned, further reducing the E/I ratio (74,75). Finally, as inhibitory circuitry reaches maturity, a new set of plasticity braking factors are triggered, including the formation of intracortical myelin and perineuronal nets, which stabilize cortical circuits and close the critical period window (6,76). Consequently, an initial decrease in the E/I ratio can signify that a critical period has been triggered and the cortex is in a relatively immature, plasticity-permissive state. As the E/I ratio reduces further, it may signify that the cortex has reached a mature, plasticity-restrictive state, with pruned excitatory synapses, fully-developed inhibitory circuitry, and mature plasticity brakes that have closed the critical period (6). As such, the greater reduction in the E/I ratio we observe in sensory systems relative to association systems may reflect that sensory systems have reached a higher degree of maturity by the end of the adolescent years while association systems remain in a more immature, plasticity-permissive state. To test this hypothesis, future work could use multimodal neuroimaging that combines our pFIC approach with other markers of critical period closure - such as intracortical myelination - to evaluate biologically-relevent signatures of when windows of critical period plasticity open and close during youth.

Our findings align with a wealth of literature demonstrating differences in the development of sensorimotor and association cortices. Studies have shown that functional connectivity, functional topography, structure-function coupling, and intrinsic dynamics follow different developmental trajectories between sensory and associative cortical systems (16,77–79). For example, while the intrinsic fluctuation amplitude of sensory systems linearly decline with age, association systems follow curvilinear developmental trajectories that peak over adolescence before declining into adulthood (80). Importantly, other recent work has shown that the development of intracortical myelin, which functions as a brake on plasticity, also varies along the sensory-to-association axis (11,81). Specifically, the period of peak growth in intracortical myelin occurs during childhood in sensorimotor cortex, yet not until adolescence in association cortex. Coupled with our current findings of a greater reduction in the E/I ratio of sensory systems (versus a weaker reduction in association systems), this work jointly indicates that sensorimotor systems are more mature by the onset of adolescence whereas association cortex may remain more plastic during the adolescent period. This interpretation aligns with a recent study showing that an fMRI marker of functional plasticity peaks during early adolescence in association cortex but continuously declines throughout childhood and adolescence in sensorimotor cortex (80).

The protracted development of the E/I ratio throughout adolescence may facilitate healthy cognitive development. We found that better cognitive ability was associated with a lower E/I ratio across the cortex in groups of age-matched youth. Since the E/I ratio normatively decreased with age, this effect may indicate that more mature cognitive performance is associated with a more mature cortical E/I ratio. As such, the E/I ratio may capture aspects of development independent of chronological age. Importantly, the magnitude of the effect was not spatially uniform. The greatest effect sizes were observed in association cortex, while the weakest effect sizes were observed in sensory cortex. This pattern is consistent with prior work showing that functional properties of the association cortex are most strongly related to cognitive performance across development (79,82). Our findings also support theoretical predictions from a biophysically-based cortical circuit model of decision-making that a balanced E/I ratio supports optimal decision-making (9). Together, our results suggest that although E/I ratio continues to develop throughout the cortex during adolescence, the development of the E/I ratio in association cortex is particularly relevant to maturing cognition. Critically, we generalized associations between the E/I ratio and cognitive ability in an independent sample of youth collected from a different continent, demonstrating the robustness of these effects across both populations and recruitment sites.

Our findings have important implications for understanding the emergence of psychopathology during adolescence. Though a prolonged period of developmental plasticity in the association cortex may be essential to healthy cognitive development, it may also represent a period of vulnerability to atypical developmental outcomes. A growing body of work has begun to implicate a disrupted E/I ratio in prefrontal cortex as a central mechanism in neuropsychiatric disorders such as depression and psychosis spectrum disorders (83–85). These conditions are thought to involve an atypically high E/I ratio in prefrontal cortex (86–89). Future studies can use our model to understand how atypical development of E/I ratio in association cortex may lead to transdiagnostic cognitive dysfunction in developmental psychopathology.

### Limitations and future work

Parameterization of local circuit parameters with the T1w/T2w ratio map and the FC gradient yielded more realistic fMRI dynamics than either gradient alone or if local circuit parameters were constrained to be spatially uniform. Future studies can explore more generic parameterizations, such as geometric eigenmodes (90).

The current study utilized parcellations with only 68 or 100 regions. Simulating the FIC model with a higher spatial resolution is computationally challenging because the number of inter-regional connections increases quadratically with the number of regions. Future work can explore more efficient algorithms. Furthermore, our analyses were limited to linear modeling of E/I ratio across a set of age bins. Future work in larger samples may facilitate the estimation of nonlinear developmental trajectories of E/I ratio.

Finally, our approach can also be used to study E/I ratio changes during cognitive tasks or during a naturalistic paradigm. When applying the pFIC model to a new dataset, dataset-specific SC, T1w/T2w ratio map and FC gradient can be used, although that might not be necessary. For example, SC, T1w/T2w ratio map and whole-cortex FC gradient were not available in the alprazolam dataset, so we utilized SC, T1w/T2w and whole-cortex FC gradient from the HCP dataset.

### Conclusion

Our results underscore the utility of large-scale circuit models to provide insights into the mechanisms driving neurocognitive development. We find that an essential aspect of healthy brain function—the cortical E/I ratio—is refined during childhood and adolescence. We also provide new evidence that this hallmark critical period mechanism is associated with improved cognitive ability. Our findings pave the way for future work to investigate how disrupted E/I balance may lead to cognitive dysfunction in psychopathology that emerges during youth and is characterized by atypical development of association cortex that undergoes protracted maturation.

## 4. Methods

We utilized the HCP S1200 release (N = 1004; **Figure 2**), pharmacological (benzodiazepine alprazolam) fMRI dataset (N = 45; **Figure 3**), PNC dataset (N = 885; **Figures 4 and 5**) and the GUSTO cohort (N = 154; **Figure 6**). In the case of HCP, we used the publicly available ICA-FIX MSMAll resting-state fMRI data in fsLR surface space. For alprazolam and PNC datasets, we used preprocessed fMRI data from our previous study, which involved slice time correction, motion correction, field distortion correction and anatomical CompCor (26). In the case of the alprazolam dataset, no resting-state fMRI was available, so we used task-fMRI after regressing out the task regressors, following our previous study (26). To be consistent, the GUSTO dataset was also preprocessed in a similar fashion as the PNC dataset. More details can be found in the Supplementary Methods.

After preprocessing, static FC was computed using Pearson’s correlation for all datasets. FCD was computed using sliding window length of ∼60 seconds, corresponding to windows of length 83, 20, 20 and 23 for the HCP, alprazolam, PNC and GUSTO datasets respectively. The window length followed best practice recommendations from previous studies (51,91). SC was computed based on the number of streamlines generated with probabilistic tractography using MRtrix3 (92). More details can be found in the Supplementary Methods.

The pFIC model (33) was fitted to the different datasets using the covariance matrix adaptation evolution strategy (CMA-ES) (56). The fitted pFIC model was used to simulate the synaptic gating variable time courses S_E_ and S_I_ of the excitatory and inhibitory populations respectively. The E/I ratio was defined as the ratio between the temporal average of S_E_ and S_I_. More details can be found in the Supplementary Methods.

The HCP data is publicly available (https://www.humanconnectome.org/). The GUSTO dataset can be obtained via a data transfer agreement (https://www.gusto.sg/). The PNC dataset is publicly available in the Database of Genotypes and Phenotypes (dbGaP accession phs000607.v3.p2). All pharmacological imaging data necessary to evaluate the conclusions in the paper are available here (https://github.com/YeoPrivateLab/CBIG_private/tree/develop/stable_projects/fMRI_dynamics/Zhang2024_pFIC/replication/Alprazolam). Code for this study can be found here (https://github.com/ThomasYeoLab/CBIG/tree/master/stable_projects/fMRI_dynamics/Zhang2024_pFIC). Co-authors (TZ and LA) reviewed the code before merging into the GitHub repository to reduce the chance of coding errors.

## Acknowledgements

Our research is supported by the NUS Yong Loo Lin School of Medicine (NUHSRO/2020/124/TMR/LOA), the Singapore National Medical Research Council (NMRC) LCG (OFLCG19May-0035), NMRC CTG-IIT (CTGIIT23jan-0001), NMRC STaR (STaR20nov-0003), Singapore Ministry of Health (MOH) Centre Grant (CG21APR1009), the Temasek Foundation (TF2223-IMH-01), and the United States National Institutes of Health (R01MH120080 & R01MH133334). Our computational work was partially performed on resources of the National Supercomputing Centre, Singapore (https://www.nscc.sg). Any opinions, findings and conclusions or recommendations expressed in this material are those of the authors and do not reflect the views of the Singapore NRF, NMRC, MOH or Temasek Foundation. In RIE2025, the GUSTO dataset was supported by funding from the NRF’s Human Health and Potential (HHP) Domain, under the Human Potential Programme. The alprazolam sample was funded by AstraZeneca Pharmaceuticals LP. DHW was also supported by NARSAD and the Sidney R. Baer, Jr. Foundation. BL was funded by NIH K99MH127293 and T32MH014654. VJS was supported by a National Science Foundation Graduate Research Fellowship (DGE-1845298). The PNC was funded via RC2 grants from the National Institute of Mental Health: MH089983 and MH089924. Additional support was provided by R01MH113550, R01MH120482, R01MH112847, R01EB022573, RF1MH116920, RF1MH121867, R37MH125829, T32MH019112, the AE Foundation, and the Penn-CHOP Lifespan Brain Institute. Data were in part provided by the Human Connectome Project, WU-Minn Consortium (Principal Investigators: David Van Essen and Kamil Ugurbil; 1U54MH091657) funded by the 16 NIH Institutes and Centers that support the NIH Blueprint for Neuroscience Research; and by the McDonnell Center for Systems Neuroscience at Washington University.

## Supplemental Material

This supplemental material consists of Supplemental Methods and Supplemental Results to complement the Methods and Results sections in the main text.

## Supplemental Methods

The HCP data collection was approved by a consortium of institutional review boards (IRBs) in the United States and Europe, led by Washington University in St Louis and the University of Minnesota (WU-Minn HCP Consortium). Data collection and study procedures for the Alprazolam dataset were approved by the University of Pennsylvania IRB; data collection for the PNC was approved by IRBs from both the University of Pennsylvania and the Children’s Hospital of Philadelphia. The GUSTO data collection was approved by the National Healthcare Group Domain Specific Review Board and the SingHealth Centralised Institutional Review Board. All participants provided informed consent before data collection. The current study was approved by the IRB of the National University of Singapore.

### S1. Human Connectome Project (HCP) dataset

We considered 1004 participants from the Human Connectome Project (HCP) S1200 release (1). All participants were scanned on a customized Siemens 3T Skyra using a multi-band sequence. Four resting-state fMRI (resting-fMRI) runs were collected for each participant in two sessions on two different days. Each resting-fMRI run was acquired with a repetition time (TR) of 0.72 s at 2 mm isotropic resolution and lasted for 14.4 min. The diffusion imaging consisted of 6 runs, each lasting ∼9 min and 50 s. Diffusion weighting consisted of 3 shells of b = 1000, 2000, and 3000 s/mm^2^ with an approximately equal number of weighting directions on each shell. Details of the data collection can be found elsewhere (1). The 1004 participants were randomly divided into training (N = 335), validation (N = 335) and test (N = 334) sets.

### S2. HCP preprocessing

Details of the HCP preprocessing can be found in the HCP S1200 manual. We utilized resting-fMRI data, which had already been projected to fsLR surface space, denoised with ICA-FIX and smoothed by 2 mm. For each run of each participant, the fMRI data were averaged within each Desikan–Killiany (2) region of interest (ROI) to generate a 68 × 1200 matrix. Each 68 × 1200 matrix was used to compute 68 × 68 FC matrix by correlating the time courses among all pairs of time courses. The FC matrices were then averaged across runs of participants within the training (or validation or test) set, resulting in a group-averaged training (or validation or test) FC matrix.

Functional connectivity dynamics (FCD) were computed as follows. We defined a window with a length of 60s (equivalent to 83 time points or TRs) as recommended by previous studies (3,4). The window was moved from the first frame to the 1118^th^ frame of BOLD time series, resulting in 1118 sliding windows in total. For each run of each participant, FC was computed within each of 1118 sliding windows. Each sliding window FC matrix was then vectorized by only considering the upper triangular entries. The vectorized FCs were correlated with each other generating a 1118 × 1118 FCD matrix. Unlike static FC, we note that the FCD matrices could not be directly averaged across participants because there was no temporal correspondence between participants during the resting-state.

In the case of diffusion MRI, probabilistic tractography was run for each participant using the fiber orientation distribution (iFOD2) algorithm provided by MRtrix3 (5). A structural connectivity (SC) matrix was generated for each participant, where each entry corresponded to the number of streamlines between two ROIs. To generate a group-level SC matrix, a thresholding procedure was employed to remove false positives. More specifically, if <50% of participants had a non-zero value in a particular entry in the SC matrix, then the entry is set to zero in all individual-level SC matrices. For each SC entry, the number of streamlines was averaged across participants with non-zero streamlines and then log-transformed (6). The entries along the main diagonal were set to 0. Group-level SCs were computed by averaging individual-level SCs within the training, validation and test sets separately and normalizing the maximum value to 0.02.

### S3. Pharmacological (benzodiazepine alprazolam) dataset

The alprazolam dataset has been previously described in detail (7). Briefly, 47 adults participated in a double-blind, placebo-controlled study using the benzodiazepine alprazolam. Each participant completed two identical experimental sessions approximately 1 week apart. In one session, participants were given a 1-mg dose of alprazolam, and in the other, they were given an identical appearing placebo. One milligram of alprazolam produces an increase in GABAergic inhibition that is considered to be clinically effective (8). The order of administration was counterbalanced across participants. Alprazolam or placebo was administered 1 hour before the fMRI acquisition so that alprazolam levels and effects were near their peak at the time of data collection (8). During both sessions, participants completed an emotion identification task lasting 10.5 min, while fMRI was acquired. Task-related fMRI results have been previously reported (7). Two participants were excluded because of missing data in at least one session, yielding a final sample of 45 participants and 90 sessions in total (ages 20.9 to 56.4; mean = 39.9, standard deviation = 12.71).

All data were collected on a Siemens Trio 3T. Blood oxygen level–dependent (BOLD) fMRI data were acquired using the following parameters: TR = 3000 ms; TE = 32 ms; flip angle = 90°; FOV = 240 mm; matrix = 128 × 128; slice thickness/gap = 2/0 mm; 30 slices; effective voxel resolution = 1.875 × 1.875 × 2 mm^3^; 210 volumes. The field of view (FOV) included temporal, inferior frontal, and visual cortices as well as subcortical structures, but excluded dorsal portions of the cerebral cortex.

The 45 participants were randomly divided into training, validation, and test sets with 15 participants each. The training-validation-test split was the same for both drug and placebo sessions.

### S4. Philadelphia Neurodevelopment Cohort (PNC) dataset

Neuroimaging data were obtained from a community-based sample of 1601 youth (ages 8.1 to 23.1; mean = 14.94; standard deviation = 3.69; male/ female = 764/837) that were part of the Philadelphia Neurodevelopmental Cohort (PNC). Data collection procedures and sample characteristics have been previously described in detail (9,10). One run of resting-fMRI data was collected per participant. Following health exclusions and rigorous quality assurance, we retained 885 participants (ages 8.2 to 23.0 at first visit; mean = 15.66; standard deviation = 3.36).

All neuroimaging data were collected on the same Siemens Trio 3T scanner as was used for the alprazolam dataset. The neuroimaging procedures and acquisition parameters have been previously described in detail (9). Briefly, BOLD fMRI was acquired using similar acquisition parameters to the alprazolam dataset: TR = 3000 ms; TE = 32 ms; flip angle = 90°; FOV = 192 × 192 mm^2^; matrix = 64 × 64; 46 slices; slice thickness/gap = 3/0 mm; effective voxel resolution = 3.0 × 3.0 × 3.0 mm^3^; 210 volumes. The key difference is that the field of view in the PNC dataset covered the whole brain (unlike the alprazolam dataset).

### S5. Alprazolam and PNC functional image processing

Details about the alprazolam and PNC datasets have been previously provided in previous studies (11). The alprazolam dataset consisted of two BOLD acquisitions per participant (drug and placebo session), which were preprocessed individually. BOLD runs were slice time–corrected and then motion-corrected. Susceptibility distortion was estimated and used to compute a corrected BOLD reference for more accurate co-registration with the anatomical reference. The BOLD reference was co-registered to the T1w reference using boundary-based registration (12). Co-registration was configured with nine degrees of freedom to account for distortions remaining in the BOLD reference. Six head motion parameters (corresponding rotation and translation parameters) were estimated before any spatiotemporal filtering. The motion-correcting transformations, field distortion correcting warp, BOLD-to-T1w transformation, and T1w-to-template (MNI) warp were concatenated and applied to the BOLD time series in a single step using antsApplyTransforms (ANTs) with Lanczos interpolation. Finally, the volumetric data was projected to fsaverage6 surface space (13).

Nuisance regression relied upon anatomical CompCor (aCompCor). aCompCor principal components were estimated after high-pass filtering the preprocessed BOLD time series (using a discrete cosine filter with 128-s cutoff). 5 CompCor components were extracted from the cerebrospinal fluid (CSF) and white matter (WM) masks. To remove task effects in the alprazolam dataset, all event conditions from the emotion identification task were modeled as 5.5-s boxcars convolved with a canonical hemodynamic response function. Each of the five emotions (fear, sad, angry, happy, and neutral) was modeled as a separate regressor. In sum, 22 regressors (6 head motion parameters and their respective temporal derivatives, top 5 aCompCor components, and 5 task regressors) were jointly regressed from the BOLD time series. The same preprocessing was performed for the PNC dataset, except that no task regressor was necessary. Overall, 17 regressors (6 head motion parameters and their respective temporal derivatives and top 5 aCompCor components) were jointly regressed from the BOLD time series.

FC and FCD of the alprazolam and PNC datasets were computed in the same manner as the HCP dataset. However, the TR was longer in the alprazolam and PNC datasets than the HCP dataset. Therefore, when computing the FCD matrices, the length of each sliding window was set to be 20 timepoints (or TRs), so that the temporal length of the window was maintained at 60s.

### S6. GUSTO dataset

To generalize our findings on the association between cognition and E/I ratio, we additionally utilized an Asian cohort, Growing Up in Singapore Towards Healthy Outcomes (GUSTO) dataset (14). We considered 389 7.5-year-old children with 1 run of resting-fMRI data and relevant cognitive scores. Participants with one or more missing cognitive scores were removed, yielding a final group of 154 participants (mean age = 7.43, std = 0.13, min age = 7.24, max age = 7.88).

All neuroimaging data were collected on a Siemens Prisma scanner. The structural data were obtained with T1w MPRAGE sequence using the following acquisition parameters: TR = 2000 ms; TE = 2.08 ms; FOV = 192 × 192 mm^2^; matrix = 192 × 192; voxel resolution = 1.0 × 1.0 × 1.0 mm^3^. The resting-fMRI data were obtained with the following acquisition parameters: TR = 2620 ms; TE = 27 ms; flip angle = 90°; FOV = 192 × 192 mm^2^; matrix = 64 × 64; 48 slices; voxel resolution = 3.0 × 3.0 × 3.0 mm^3^; 120 volumes. The field of view of the GUSTO dataset covered the whole brain.

### S7. GUSTO preprocessing

For each resting-fMRI run, the following sequence of preprocessing steps were performed. The first 4 frames of the run were removed. The run was then slice time-corrected and then motion-corrected. Frames with FD > 0.5mm and DVARS > 80 were marked as motion outliers. Next, the run was co-registered to the structural image with boundary-based registration (12). Nuisance regression was performed with the inclusion of white matter and CSF signals, 6 head motion parameters and their respective temporal derivates as well as the top 5 aCompCor components (19 regressors in total were jointly applied). The motion outlier frames were then censored and interpolated. Finally, the run was bandpass-filtered (0.009Hz < f < 0.08Hz) and projected to Freesurfer fsaverage6 surface space.

FC and FCD of the GUSTO dataset were computed in the same manner as the HCP dataset. However, the TR was longer in the GUSTO than the HCP dataset. Therefore, when computing the FCD matrices, the length of each sliding window was set to be 23 timepoints (or TRs), so that the temporal length of the window was maintained at around 60s.

### S8. Feedback Inhibition Control (FIC) model

The derivation of the FIC model was thoroughly described in a previous study (15). Here we provide some intuition for the FIC model. The neuronal activities of the *j*-th cortical region follow the nonlinear differential equations shown below

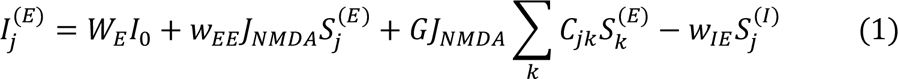

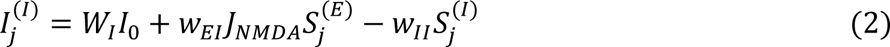

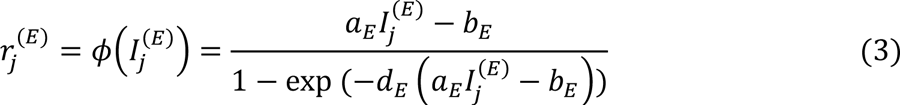

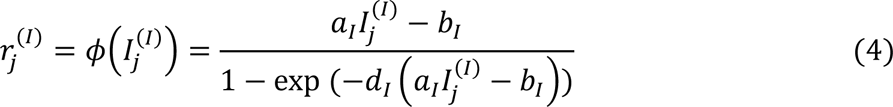

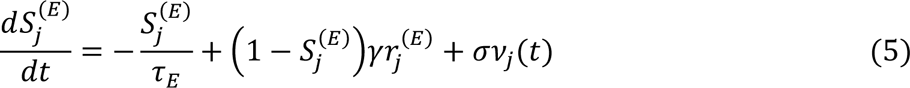

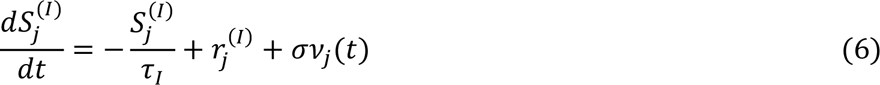

where *S*, *r*, and *I* represent synaptic gating variables, firing rate, and synaptic currents respectively. The superscripts *E* and *I* denote the excitatory and inhibitory neuronal populations respectively.

The input current 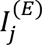 of the excitatory population of the *j*-th cortical ROI is the sum of four inputs (Equation 1). The first input is the external input current *W*_*E*_ *I*_0_, which might include subcortical delays. The second input is the intra-regional excitatory-to-excitatory current governed by the excitatory-to-excitatory recurrent connection strength *w*_*EE*_ scaled by the synaptic coupling constant *J_NMDA_*. The third input is the inter-regional input, which is controlled by the SC matrix (*C*_*jk*_ is the connectivity between regions *j* and *k*) and scaled by the global constant *G*. The fourth input is the intra-regional negative feedback from the inhibitory population governed by the inhibitory-to-excitatory connection strength *w*_*IE*_.

The input current 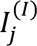 of the inhibitory population of the *j*-th cortical ROI is the sum of three inputs (Equation 2). The first input is the external input current *W*_*I*_*I*_0_. The second input is the intra-regional positive feedback from the excitatory population governed by the excitatory-to-inhibitory connection strength *w_EI_* scaled by the synaptic coupling constant *J*_*NMDA*_. The third input is the intra-regional inhibitory-to-inhibitory current governed by the inhibitory-to-inhibitory recurrent connection strength *w*_*II*_.

The excitatory input current 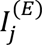 and inhibitory input current 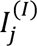 are transformed into firing rates via the input-output functions specified in Equations 3 and 4. Following previous studies (15), the parameters of the input-output function were set to be *a*_*E*_ = 310n/C, *a*_*I*_ = 615n/C, *b*_*E*_ = 125Hz, *b*_*I*_ = 177Hz, *d*_*E*_ = 0.16s and *d*_*I*_ = 0.087s. Finally, the rate of change of the synaptic gating variables 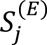 and 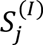 are computed via equations 5 and 6. Following previous studies (15), the kinetic parameters for synaptic activities *τ*_*E*_, *τ*_*I*_ and *γ* were set to 100ms, 10ms and 0.641 respectively. *v*_*j*_ (*t*) corresponds to uncorrelated standard Gaussian noise with the noise amplitude being controlled by *σ*.

Following the original study (15), *w*_*II*_, *W*_*E*_, *W*_*I*_, *I*_0_ and *J*_*NMDA*_ were set to 1, 1, 0.7 0.382nA and 0.15nA respectively in the current study. The inhibitory-to-excitatory connection strength *w*_*IE*_ was computed analytically to ensure that the excitatory firing rate is maintained to be around 3Hz (16). We note that this analytical computation assumes a noiseless system, so in practice, we imposed a constraint that the firing rate is between 2.7Hz to 3.3Hz. During the estimation of the pFIC model (next section), parameters were rejected if firing rates fall outside this range.

The excitatory-to-excitatory recurrent connection strength *w*_*EE*_, excitatory-to-inhibitory connection strength *w*_*EI*_, noise amplitude *σ* and global SC scaling constant *G* are unknown parameters, which will be estimated by fitting to empirical fMRI data (next section). Given a fixed set of model parameters, equations 1 to 6 can be used to simulate the time courses of excitatory and inhibitory synaptic gating variables 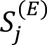 and 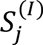 of each ROI. The regional E/I ratio was defined as the ratio between the temporal average of 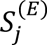 and 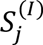. The mean cortical E/I ratio was the average of regional E/I ratios across all cortical ROIs. The simulated excitatory synaptic gating variables (*S*^(*E*)^) were also fed to the Balloon-Windkessel hemodynamic model to simulate fMRI BOLD signals (15, 17,18). The simulated fMRI BOLD signals were then used to generate simulated static FC and FCD.

### S9. Parametric FIC (pFIC) model

Recall that the FIC model was instantiated using the Desikan-Killiany parcellation with 68 regions of interest. Given that we wanted the excitatory-to-excitatory recurrent connection strength w_EE_, excitatory-to-inhibitory connection strength w_EI_ and noise amplitude *σ* to be spatially heterogeneous (with *G* being a global constant), if we optimized each parameter independently, there would be a total of 68 × 3 + 1 = 205 parameters, which is computationally challenging.

In our previous study (19), we reduced the number of “free” parameters by parameterizing the local synaptic parameters with a linear combination of the first principal FC gradient and T1w/T2w ratio map. We note that in the previous study (19), we considered a highly simplified parametric mean field model that does not differentiate between excitatory and inhibitory neural populations. Here, we considered the same approach for the FIC model by parameterizing w_EE_, w_EI_, and *σ* as a linear combination of T1w/T2w myelin map and first principal FC gradient.

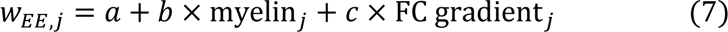

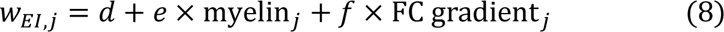

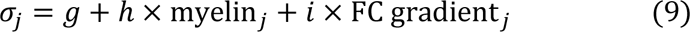

where *j* denotes the ROI index. By adopting this parameterization approach, the number of “free” numbers was reduced to 3 × 3 + 1 = 10 parameters.

### S10. Optimization of the parametric FIC (pFIC) model in the HCP dataset

The 10 unknown parameters of the pFIC model were optimized using a previously published approach (19) by maximizing fit to empirical static FC and FCD. The agreement between empirical and simulated FC matrices was defined as the Pearson’s correlation (*r*) between the z-transformed upper triangular entries of the two matrices. Larger *r* indicates more similar static FC. However, Pearson’s correlation does not account for scale difference, so we also computed the absolute difference (*d*) between the means of the empirical and simulated FC matrices (16). A smaller d indicates more similar static FC.

We note that there is no temporal correspondence between simulated and empirical FCD matrices, so we cannot simply use the Euclidean distance to measure dissimilarity. Instead, the dissimilarity between simulated and empirical FCD matrices was quantified by using the Kolmogorov-Smirnov (*KS*) distance. Here, the KS distance was defined as the maximum distance between the cumulative distribution functions (CDFs) constructed by collapsing the upper triangular entries of simulated and empirical FCD matrices (19,20). Hence, a small *KS* distance indicated 2 similar CDFs, thepre 2 similar FCD matrices. Because the *KS* distance was computed by collapsing the upper triangular entries of the FCD matrices, no temporal correspondence was assumed.

The overall cost was defined as (1 – *r*) + *d* + *KS*. A smaller cost indicates better agreement between simulated and empirical fMRI. Recall that the 1004 HCP participants were randomly divided into training (N = 335), validation (N = 335) and test (N = 334) sets. Following our previous study (19), we used the covariance matrix adaptation evolution strategy (CMA-ES) (21) to minimize the overall cost function during training. Due to the lack of temporal correspondence between FCD matrices across runs and participants during rs-fMRI scans, directly averaging FCD matrices will cancel out the temporal dynamics. Representing FCD matrices using CDFs avoided such problem because averaging the CDFs across runs and participants could still largely preserve the distribution of FCD matrix entries. We thus averaged the FCD CDFs across participants and runs separately within the training, validation and test sets.

In the HCP training set, the CMA-ES algorithm was iterated 100 times and repeated 5 times with different random initializations, yielding a total of 500 candidate parameter sets (**Figure 2A**). The 500 candidate parameter sets were evaluated in the validation set to obtain the top 10 candidate parameter sets. To ensure diversity among the parameter sets, the procedure to select the top 10 parameter sets was as follows. First, the parameter set with the lowest validation cost was selected. Then, the parameter set with the lowest validation cost and whose parameter (*w_EE_*, *w_EI_*, *σ*) maps exhibited less than 0.98 correlation with the current selected parameter set(s) was selected. This procedure was repeated until 10 parameter sets were selected.

The top 10 candidate parameter sets from CMA-ES were then applied to the HCP test set SC. For each parameter set, 1000 simulations were performed, yielding 1000 simulated static FC and FCD matrices. Pearson correlation and the absolute difference were then computed between each simulated FC and the empirical FC from the HCP test set and averaged. Similarly, KS statistics was computed between each simulated FCD CDF and the empirical FCD CDF from the HCP test set and averaged.

To speed up the computation, a step size of 6ms was used to integrate the ODEs in the training set. However, to ensure more accurate integration, a step size of 0.5ms was used for both validation and test sets. To ensure that this time step size was small enough, we repeated the experiment using a step size of 6ms for the training set and a step size of 0.1ms for the validation and test sets. The overall cost was highly similar across step size of 0.5ms and step size of 0.1ms. In particular, for both 0.5ms and 0.1ms step sizes, an overall cost of 0.58 ± 0.018 was achieved in the HCP test set across the top 10 parameter sets from the validation set.

Additionally, we observed that the correlations between w_EI_ and T1w/T2w ratio were consistently negative, while the correlations between w_EI_ and RSFC gradient were consistently positive across the top 10 parameter sets. Since the training sets for the alprazolam and PNC datasets were substantially smaller than the HCP training set (∼15 participants versus 335 participants), when optimizing the pFIC model in the alprazolam, PNC and GUSTO datasets, we additionally imposed the constraints that w_EI_ and T1w/T2w ratio should be negative, while the correlations between w_EI_ and FC gradient should be positive.

It is worth noting that when evaluating the top 10 model estimates (selected from the HCP validation set) in the HCP test set, the correlation loss (1-*r*) ranged from 0.27 to 0.29, absolute difference loss *d* ranged from 0.08 to 0.14, and the *KS* distance ranged from 0.14 to 0.23. In our previous study (19), our cost function was (1-*r*) + *KS*. Changing the relative weights of (1-*r*) and *KS* did not substantially change the model estimate. Thepre, we kept the weights unchanged in the current study. However, we observed that the simulated fMRI time courses were overly synchronized, so we included the additional absolute difference (*d*) metric in the current study. We observed that setting the relative weight of *d* to be the same as the other 2 terms was sufficient to prevent the simulated fMRI time courses from becoming over-synchronized (**Figure S1**). Thepre, we did not consider changing the relative weights further.

### S11. Pharmacological E/I ratio analysis

Recall that the 45 participants of the alprazolam dataset were randomly divided into training (N = 15), validation (N = 15) and test (N = 15) sets. Each participant had 2 runs of fMRI data - one run for the drug session and one run for the placebo session. Because there was no diffusion data in the alprazolam dataset, the SC matrices used for training, validation, and testing were the same as HCP training, validation, and tests set respectively. The T1w/T2w ratio map also had to be generated from HCP training set.

Due to the limited FOV of the alprazolam dataset, we only considered regions with more than 50% ROI coverage, resulting in 42 ROIs (in the case of the Desikan-Killiany parcellation). The remaining 26 ROIs were masked out for the group-level SC, static FC, FCD, T1w/T2w ratio. Furthermore, given the limited FOV, the first principal FC gradient was generated from HCP training set and the 26 ROIs were masked out similarly. For each experimental condition (placebo or alprazolam), 250 candidate parameter sets were generated from the condition’s training set. The top 10 parameter sets from the validation set were evaluated in the test set.

Although the parameters were only estimated based on the 42 ROIs, the estimated linear coefficients could be used to generate whole cortex estimates of w_EE_, w_EI_ and *σ* (based on equations 7 to 9) given that the original T1w/T2w ratio and FC gradient (from the HCP training set) covered the whole cortex. The simulated time courses S_E_ and S_I_ were then generated using the 68-ROI SC and extrapolated model parameters. For a given set of parameters, 1000 simulations were performed to generate 1000 sets of E/I ratio. The final E/I ratio was the average across 1000 sets of E/I ratio.

E/I ratio contrast was defined as the difference between E/I ratio of the placebo sessions and E/I ratio of the drug sessions. To test that the E/I ratio contrast was significantly greater than 0, we performed permutation test to generate a null distribution of E/I ratio contrasts. More specifically, after dividing the participants into training, validation and test sets, the ‘drug’ and ‘placebo’ sessions were randomly permuted within each participant. The entire procedure (above) was repeated, generating a null value for the E/I ratio contrast. The permutation procedure was repeated 100 times, yielding a null distribution of regional E/I ratio contrasts. A 2-tail p-value was computed based on this null distribution.

The regional E/I ratio contrast was also correlated with benzodiazepine receptor (BZR) density. The statistical significance of this correspondence was computed using a spin test that accounts for spatial autocorrelation (22).

### S12. Association between age and E/I ratio in the PNC dataset

885 participants of the PNC dataset (9) were first sorted according to age in ascending order and divided into 29 groups of 30 or 31 participants. For each age group, 15 participants were randomly selected as the validation set, while the remaining participants were assigned to the training set.

To be consistent with previous analyses, the SC matrices used for training and validation were the same as HCP training and validation sets respectively. Both T1w/T2w ratio and the first principal FC gradient maps were generated from HCP training set, consistent with the alprazolam analyses. For each age group, 250 candidate model parameter sets were generated from the group’s training set using CMA-ES and evaluated in the group’s validation set. For each age group, the parameter set with the lowest validation cost was used to estimate regional E/I ratio across the cortex.

For each cortical ROI, we fitted a linear regression model between regional E/I ratio and the mean age of each age group. The slope of the linear regression model (for each brain region) was visualized on the cortical surface. All slopes were negative and all p-values survived FDR correction (q < 0.05). For robustness, the split of the participants into training and validation sets were repeated 5 times and the most representative split was shown in **Figure 4**.

### S13. Association between cognition and E/I ratio in the PNC dataset

Each PNC participant had completed a set of 12 tasks from 4 cognitive domains, including executive control, episodic memory, complex cognition, and social cognition. Three types of scores, including an accuracy score, a speed score, and an efficiency score, were obtained for each of the 12 tasks. Factor analyses of each type of scores were performed within each cognitive domain and across all cognitive domains to generate three domain-specific accuracy factor scores and one domain-general (overall) accuracy score (23).

To control for age, 885 participants were sorted according to age in an ascending order. For each 12-month interval, participants whose age were within this interval were extracted to form one age group. For each age group, participants with domain-general (overall) accuracy scores above the median were assigned to a high-performance group, the rest are assigned to a low-performance group. In sum, 441 participants were assigned to the high-performance group (mean age = 15.68), 444 participants were assigned to the low-performance group (mean age = 15.64). Both high and low-performance groups were then divided into 14 subgroups of 31 (or 32) participants, yielding 14 pairs of age-matched high-performance and low-performance groups (see **Figure 5A and 5B**).

Participants of each subgroup were further randomly divided into a training set (N = 16) and a validation set (N = 15 or 16). Similar to the previous analyses, SCs used for training and validation were from the HCP training and validation sets respectively. T1w/T2w ratio and FC gradient maps were from the HCP training set. The E/I ratios of the 14 pairs of high-performance and low-performance groups were estimated separately and compared using a 2-tail 1-sample t-test. To test for domain specificity, we also repeated the analyses for the 3 domain-specific accuracy scores. FDR correction with *q* < 0.05 was used to correct for multiple comparisons.

### S14. Association between cognition and E/I ratio in the GUSTO dataset

Our analyses were based on the fMRI and behavioral data of 154 participants from the GUSTO dataset. The fMRI data were acquired when the participants were around 7.5 years old. We selected 5 behavioral scores which assessed participants’ cognitive performances. All behavior tests were performed within 1.5 years of fMRI acquisition (i.e., age 6 to 8.5). The 5 test scores were as follows: (1) Cambridge Neuropsychological Test Automated Battery (CANTAB) Spatial Working Memory (SWM) test (completed at age 6): sum of total errors for 4 and 6 boxes trails. (2) Delayed Matching to Sample (DMS) test (completed at age 6): percentage of the total number of trials upon which a correct selection was made on the participant’s first response. (3) Behavior Rating Inventory of Executive Function (BRIEF; completed at age 7): Cognition Regulation Index T-score. (4) Wechsler Abbreviated Scale of Intelligence (WASI) test (completed at age 7): sum of Block Design and Matrix Reasoning T-scores. (5) CANTAB SWM test (completed at age 8.5): sum of total errors for 4 and 8 boxes trails.

Principal component analysis (PCA) was performed on these 5 scores across all participants to derive the first principal component (PC1) score. Higher PC1 score indicated better cognitive performance across the 5 behavioral scores (on average). Participants were sorted according to their PC1 scores in an ascending order. The first 77 participants were assigned to the low-performance group, while the rest of 77 participants were assigned to the high-performance group. Ages were well-matched between high and low-performance groups (**Figure 6A**).

Participants of high and low-performance groups were further randomly divided into a training set (N = 39) and a validation set (N = 38). Similar to the previous analyses, SCs used for training and validation were from the HCP training and validation sets respectively. T1w/T2w ratio and FC gradient maps were from the HCP training set. For robustness, the analyses were repeated 5 times with different random training-validation splits of participants within high-performance group and each low-performance group. Results from the most representative split were shown in the results.

To compute the statistical significance of E/I ratio differences between low and high-performance groups, PC1 scores were randomly permuted across participants. The participants were again assigned to high or low-performance groups according to their permuted PC1 scores. Then the E/I ratio difference was re-estimated. This permutation process was repeated 100 times to construct a null distribution of E/I ratio difference. A 2-tail p-value was computed based on this null distribution.

## Supplemental Figures

**Figure S1.**
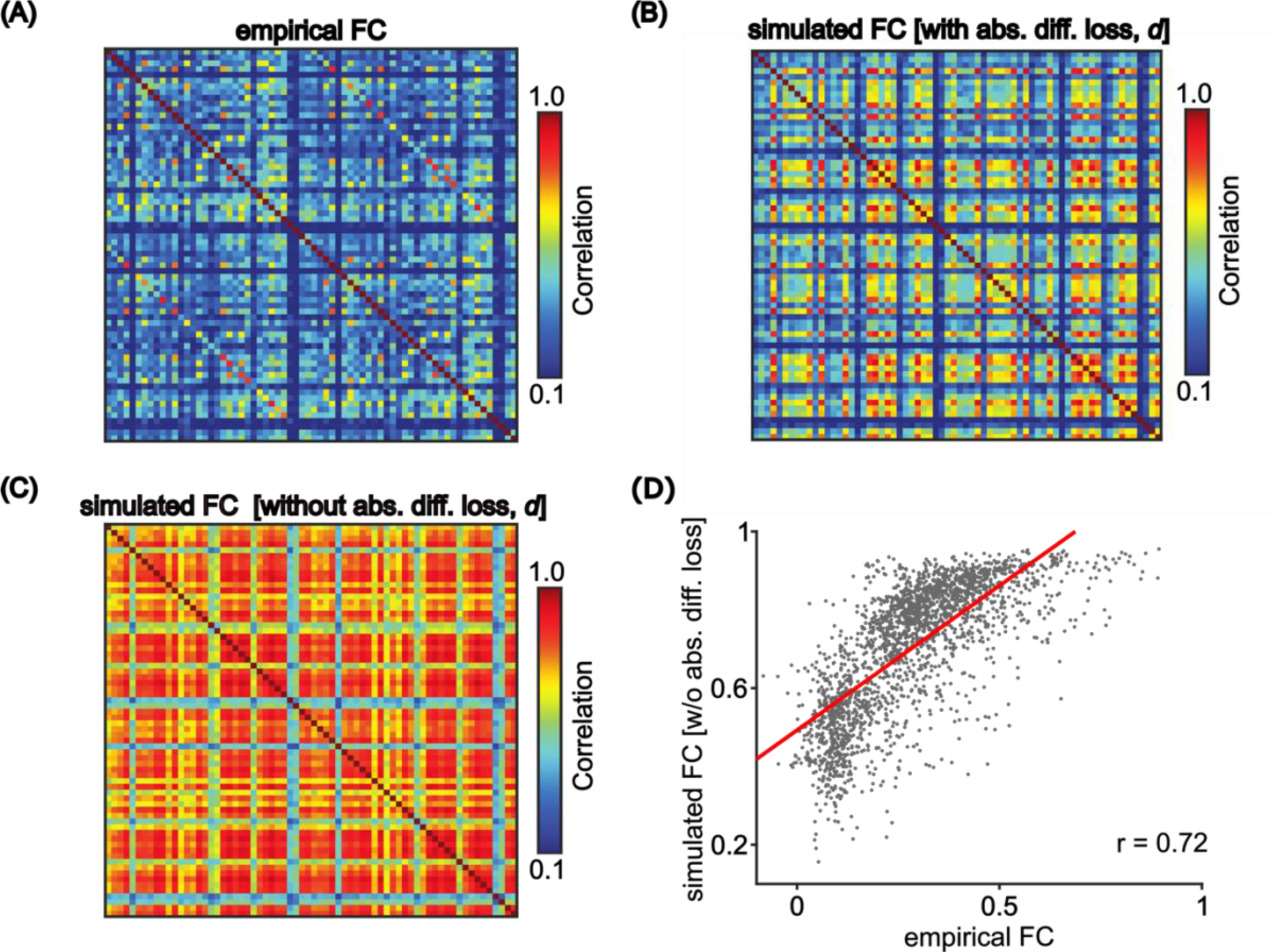
Importance of absolute difference (*d*) metric to prevent overly synchronized simulated fMRI time series. (A) Empirical FC from the HCP test set. (B) Simulated FC from the pFIC model using the best model parameters (from the validation set) and SC from the test set when the cost function contains all three terms: (1 – *r*) + *d* + *KS*. The cost function contained two terms related to static FC: disagreement between empirical and simulated FC in terms of Pearson’s correlation (1 – *r*) and absolute difference (*d*). (C) Simulated FC from the pFIC model using the best model parameters (from the validation set) and SC from the test set when the cost function contained only two terms: (1 – *r*) + *KS.* (D) Agreement (Pearson’s correlation) between empirical and simulated static FC when the when the cost function contained only two terms: (1 – *r*) + *KS*. Thepre, without the inclusion of the absolute difference (*d*) metric, we can obtain good correlation agreement between simulated and empirical FC. However, by comparing panels (B) and (C), we observe that the lack of the absolute difference (*d*) metric leads to overly synchronized fMRI signals, compared with the empirical FC in panel A. It is also worth noting that the over-synchronization phenomenon was also observed in our previous study (19), which used (1 – *r*) + *KS* as the cost function. Thepre, in this study, we added the absolute difference (*d*) cost to the cost function.

**Figure S2.**
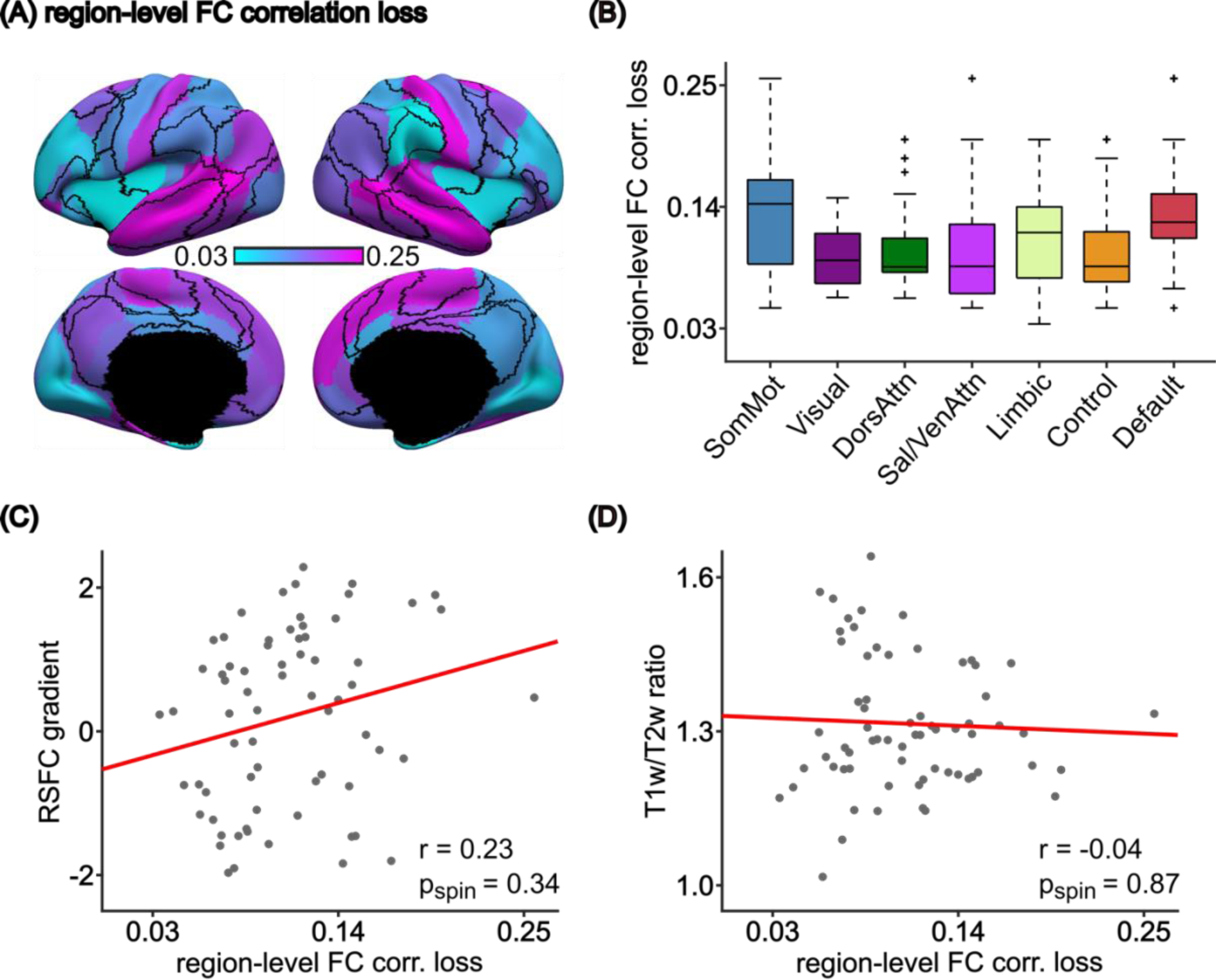
Regional evaluation of pFIC model for static FC. (A) Spatial distribution of region-level FC loss (dissimilarity) between simulated and empirical fMRI in the HCP test set. The regional FC correlation loss was defined as (1 - *r_i_*), where *r_i_* is the Pearson’s correlation between the *i*-th rows of empirical and simulated static FC matrices. (B) Regional FC correlation losses grouped by different large-scale networks. The boxes show the inter-quartile range (IQR) and the median. Whiskers indicate 1.5 IQR. Black crosses represent outliers (C) There was no significant correlation between RSFC gradient and the regional FC correlation loss (*r* = 0.23, two-tail spin test *p* = 0.34). (D) There was no significant correlation between T1w/T2w ratio and the regional FC correlation loss (*r* = −0.04, two-tail spin test *p* = 0.87).

**Figure S3.**
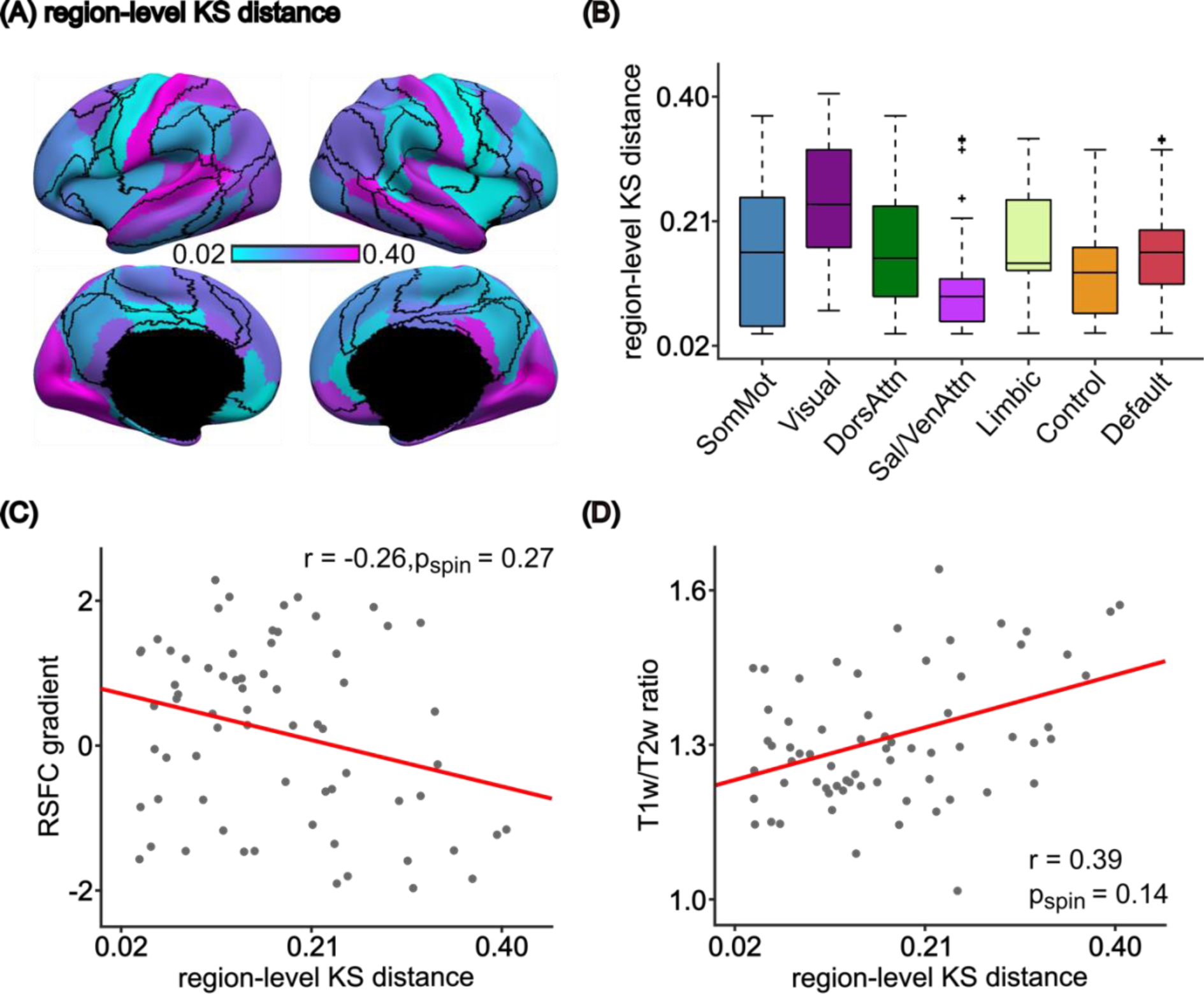
Regional evaluation of pFIC model for FCD. (A) Spatial distribution of region-level FCD dissimilarity (KS distance) between empirical and simulated FCD in the HCP test set. To compute regional KS distance, recall that we have previously computed a 68 × 68 FC matrix for each sliding window (1118 sliding windows in total). For each region, the corresponding rows of the 68 × 68 FC matrices were then correlated across the 1118 windows, yielding a 1118 × 1118 FCD matrix for each region. The KS distance can thus be computed for each region. (B) Regional KS distances grouped by different networks. The boxes show the inter-quartile range (IQR) and the median. Whiskers indicate 1.5 IQR. Black crosses represent outliers. (C) There was no significant correlation between RSFC gradient and regional KS distance (*r* = −0.26, two-tail spin test *p* = 0.27). (D) There was no significant correlation between T1w/T2w ratio and regional KS distance (*r* = 0.39, two-tail spin test *p* = 0.14).

**Figure S4.**
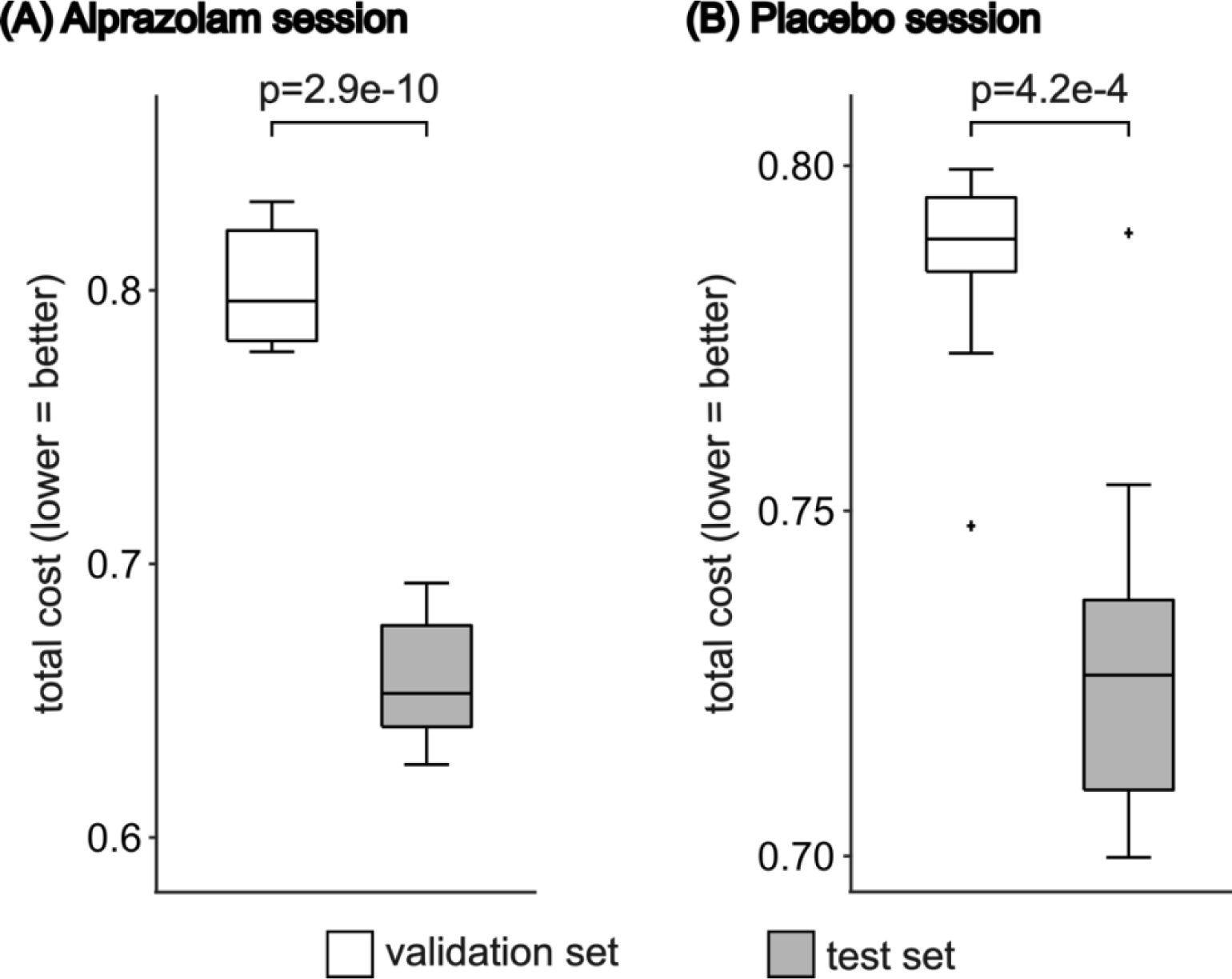
(A) Comparison of the total costs in the validation set and the test set of the alprazolam session. (B) Comparison of the total costs in the validation set and the test set of the placebo session. The total costs in the validation set correspond to the lowest 10 validation costs, generated from the top 10 sets of parameters from the validation set. The total costs of the test set were computed using the same 10 sets of parameters and the structural connectivity of the test set. For both alprazolam and placebo sessions, total costs of the test set were significantly lower than those of the validation set, suggesting that the parameters generalized well from the validation set to the test set. The boxes show the inter-quartile range (IQR) and the median. Whiskers indicate 1.5 IQR. Black crosses represent outliers.

**Figure S5.**
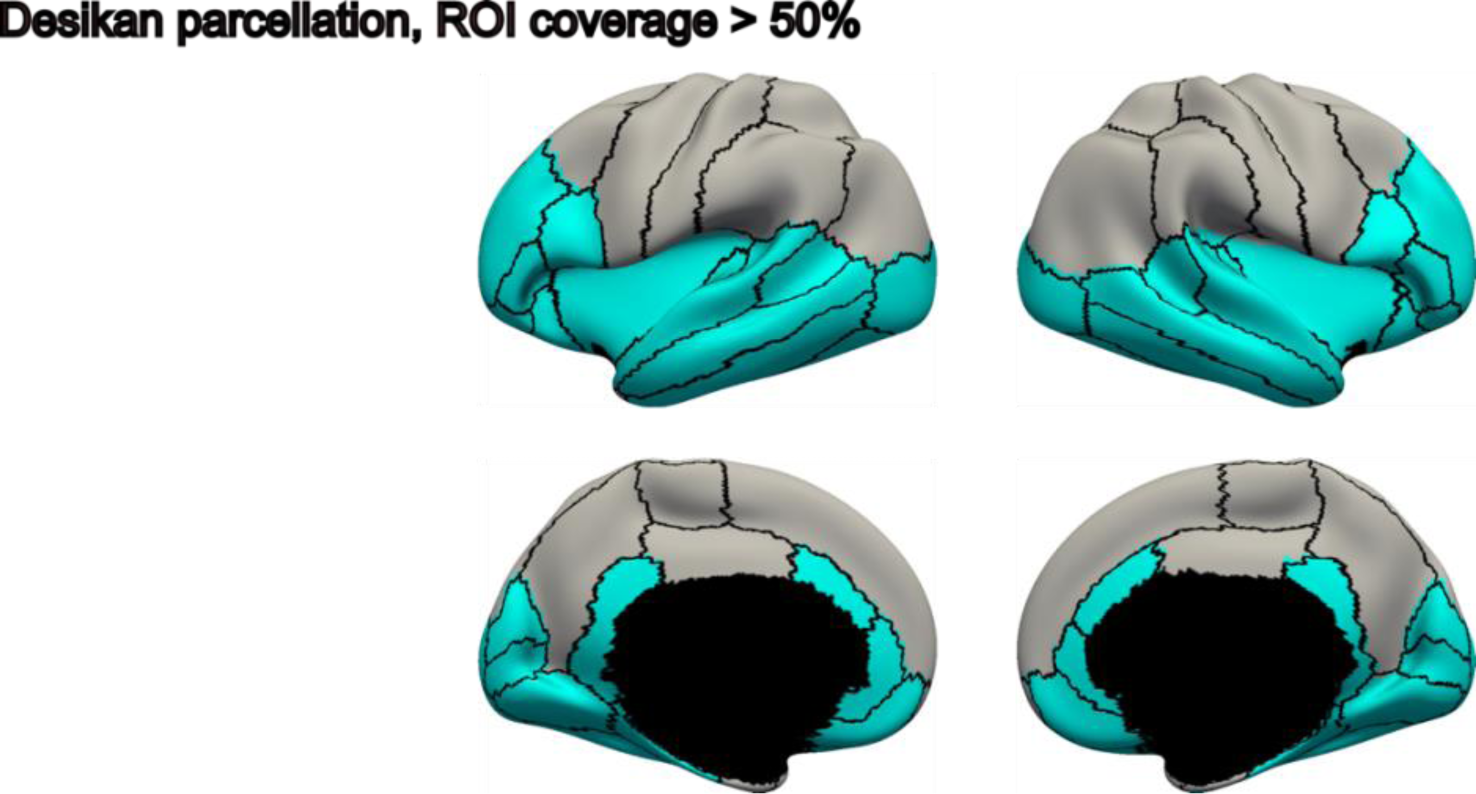
Desikan parcellation with 50% ROI coverage threshold. Due to limited FOV of the alprazolam dataset, only ROIs with coverage higher than a pre-specified threshold were included. ROIs included (excluded) for analysis are colored in cyan (grey). 42 out of 68 ROIs are included.

**Figure S6.**
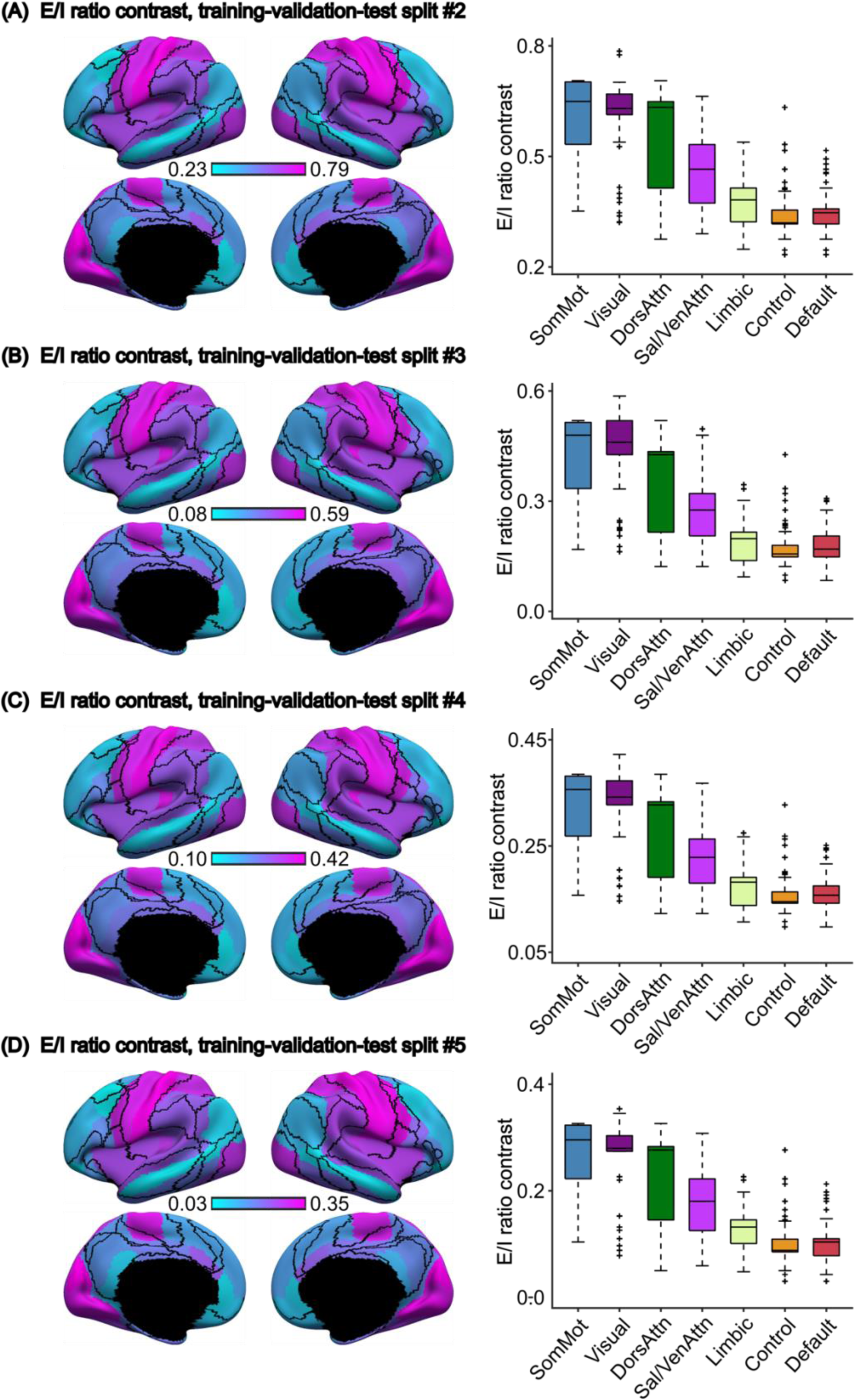
(A to D) As a control analysis of the alprazolam E/I ratio contrast result, the same analysis was replicated with 4 additional training-validation-test participant splits. E/I ratio contrast was defined as the E/I ratio difference between the placebo session and the drug session. 45 participants were randomly assigned to a different training, validation, and test set. (Left) Spatial distribution of E/I ratio contrast between placebo and drug session. (Right) The E/I ratio contrast decreases along a sensory-to-association axis. The boxes show the inter-quartile range (IQR) and the median. Whiskers indicate 1.5 IQR. Black crosses represent outliers.

**Figure S7.**
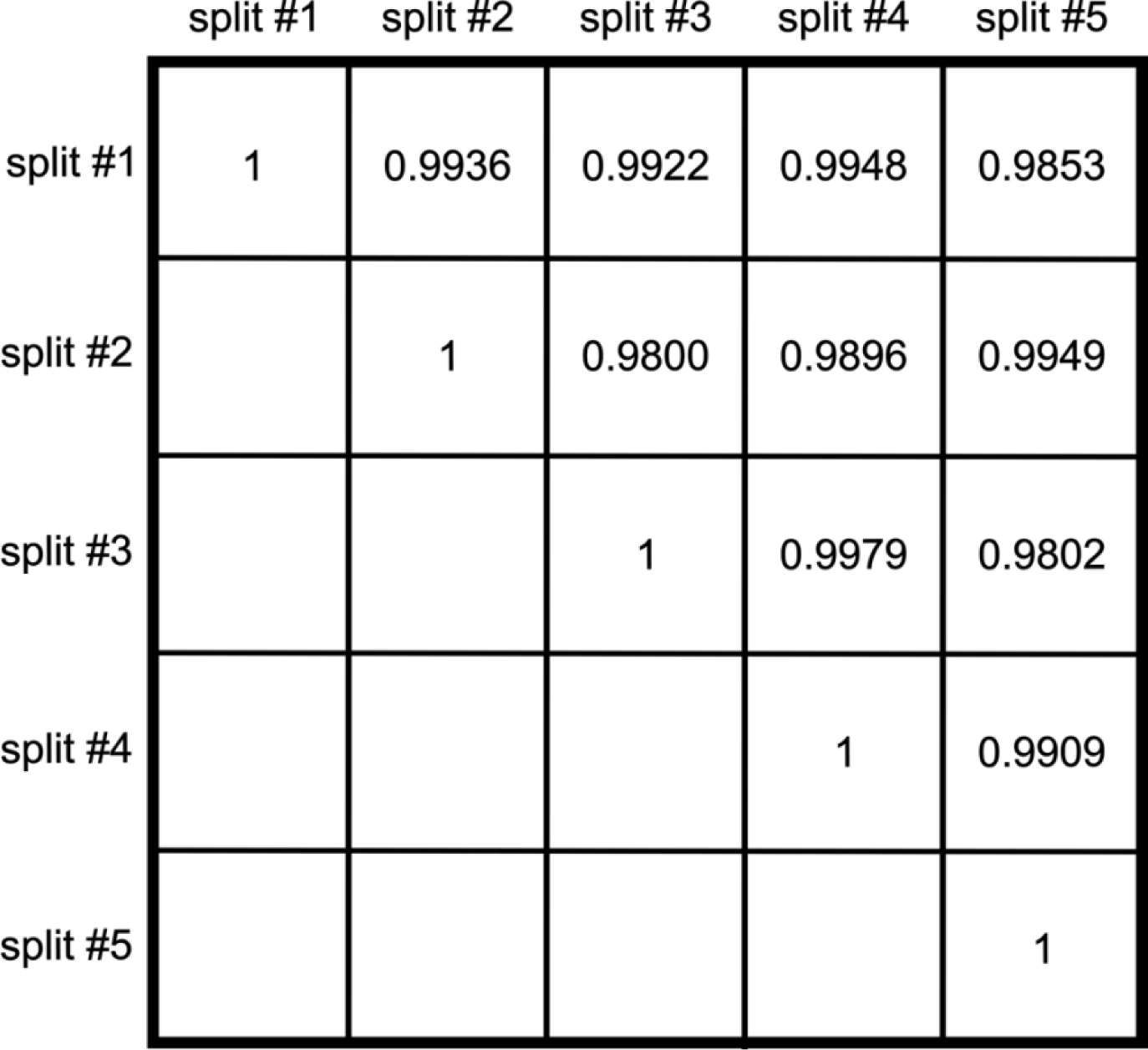
Pairwise correlation between the regional E/I ratio contrast across different training-validation-test participant splits (split #1-5). The spatial distribution of region E/I ratio contrast are highly similar across splits (*r* = 0.9899 ± 0.0062, mean ± std). Only the upper triangle of the matrix is shown. We chose the split that had the highest median correlation of regional E/I ratio contrast with the other 4 splits as the most representative split (i.e., split #1) and showed as Figure 3 of the main text.

**Figure S8.**
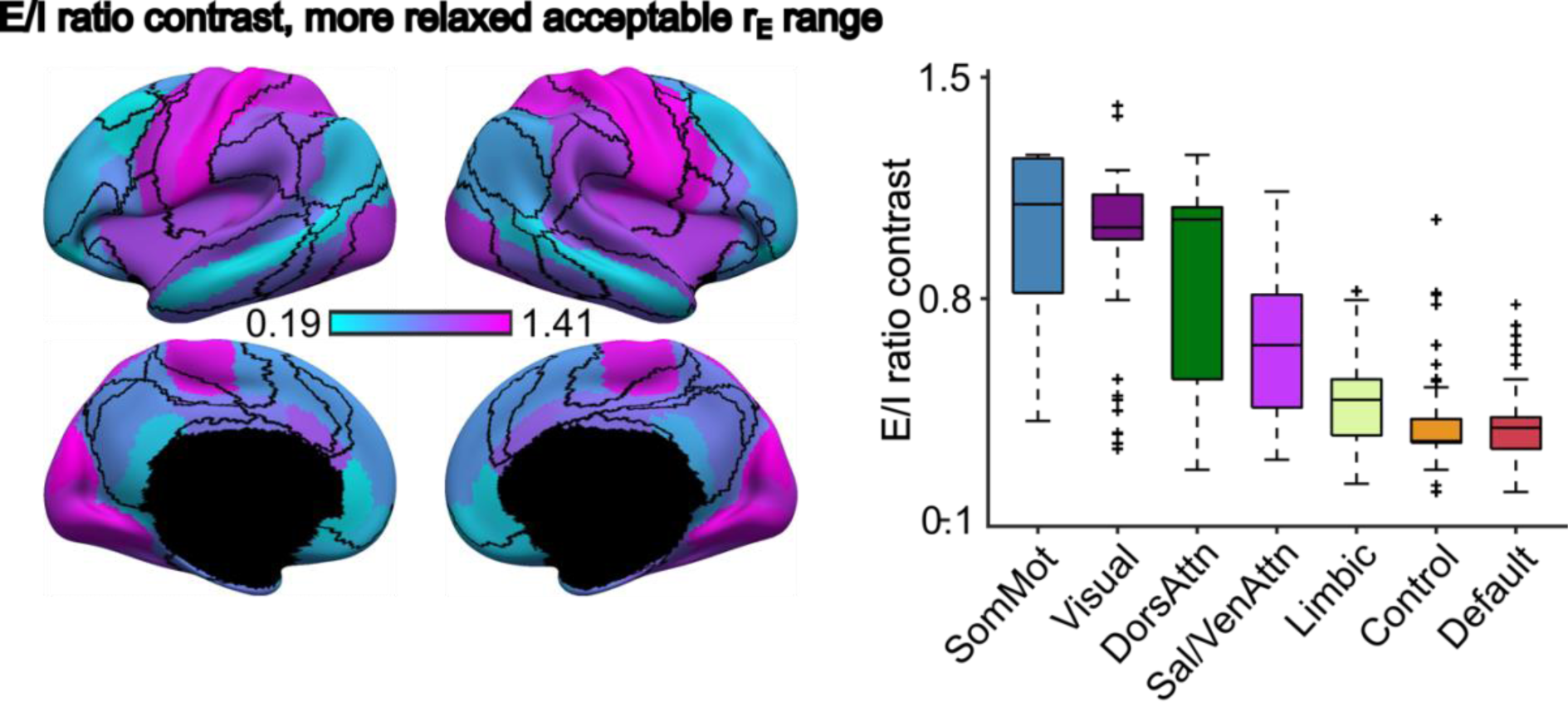
As a control analysis of the alprazolam E/I ratio contrast result, the acceptable excitatory firing rate range was set to be less strict. The range was changed from 2.7 – 3.3Hz to 2.5 – 3.5 Hz. We repeated the same analysis, and the model parameters were retrained using the same optimization approach. The boxes show the inter-quartile range (IQR) and the median. Whiskers indicate 1.5 IQR. Black crosses represent outliers.

**Figure S9.**
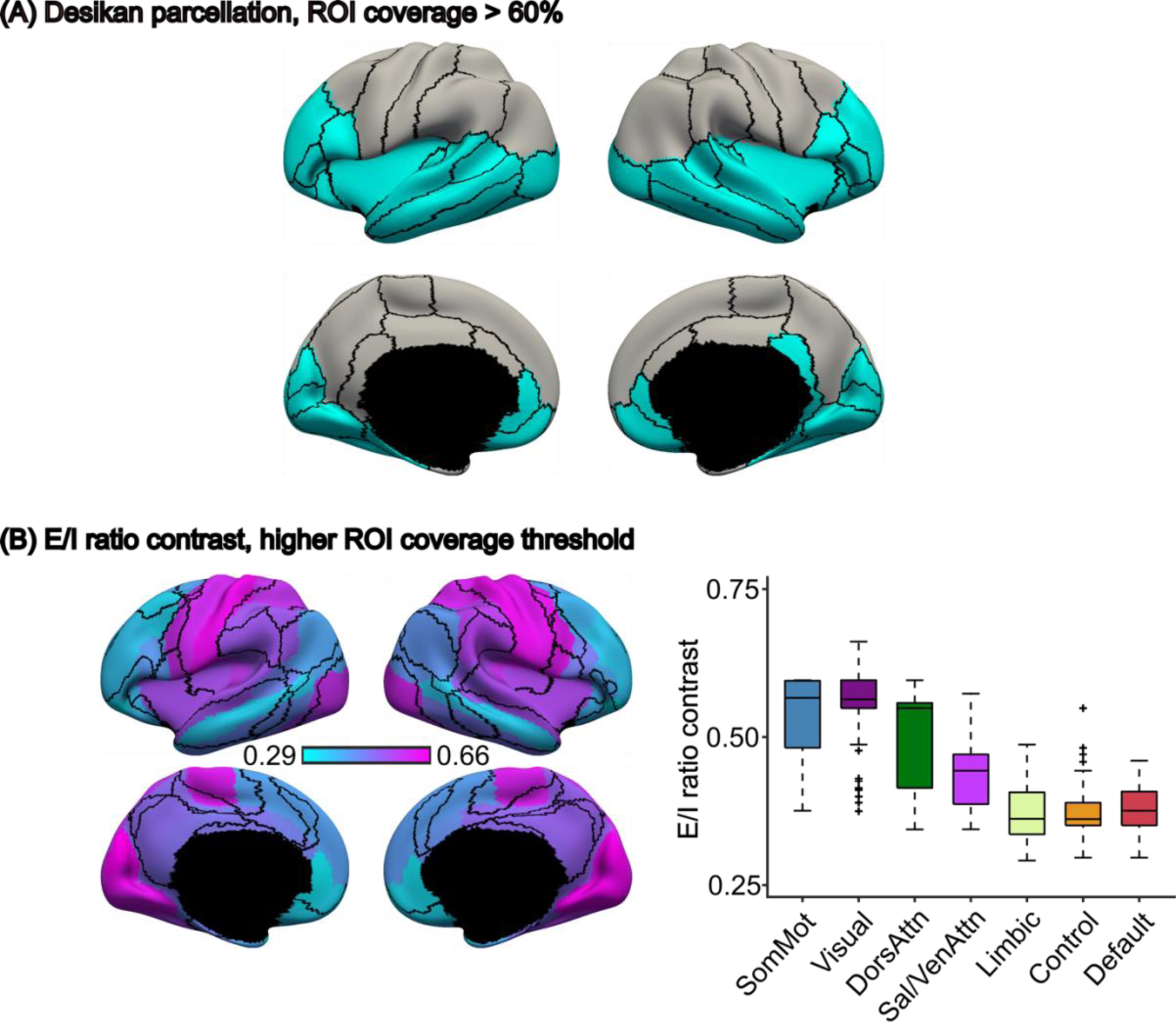
(A) Desikan parcellation with 60% ROI coverage threshold, 39 out of 68 ROIs are left. (B) As a control analysis of the alprazolam E/I ratio contrast result, the ROI coverage threshold was changed to be stricter. The threshold was raised from 50% to 60%. As a result, 3 more ROIs were removed from analysis. We repeated the same analysis, and the model parameters were retrained using the same optimization approach. The boxes show the inter-quartile range (IQR) and the median. Whiskers indicate 1.5 IQR. Black crosses represent outliers.

**Figure S10.**
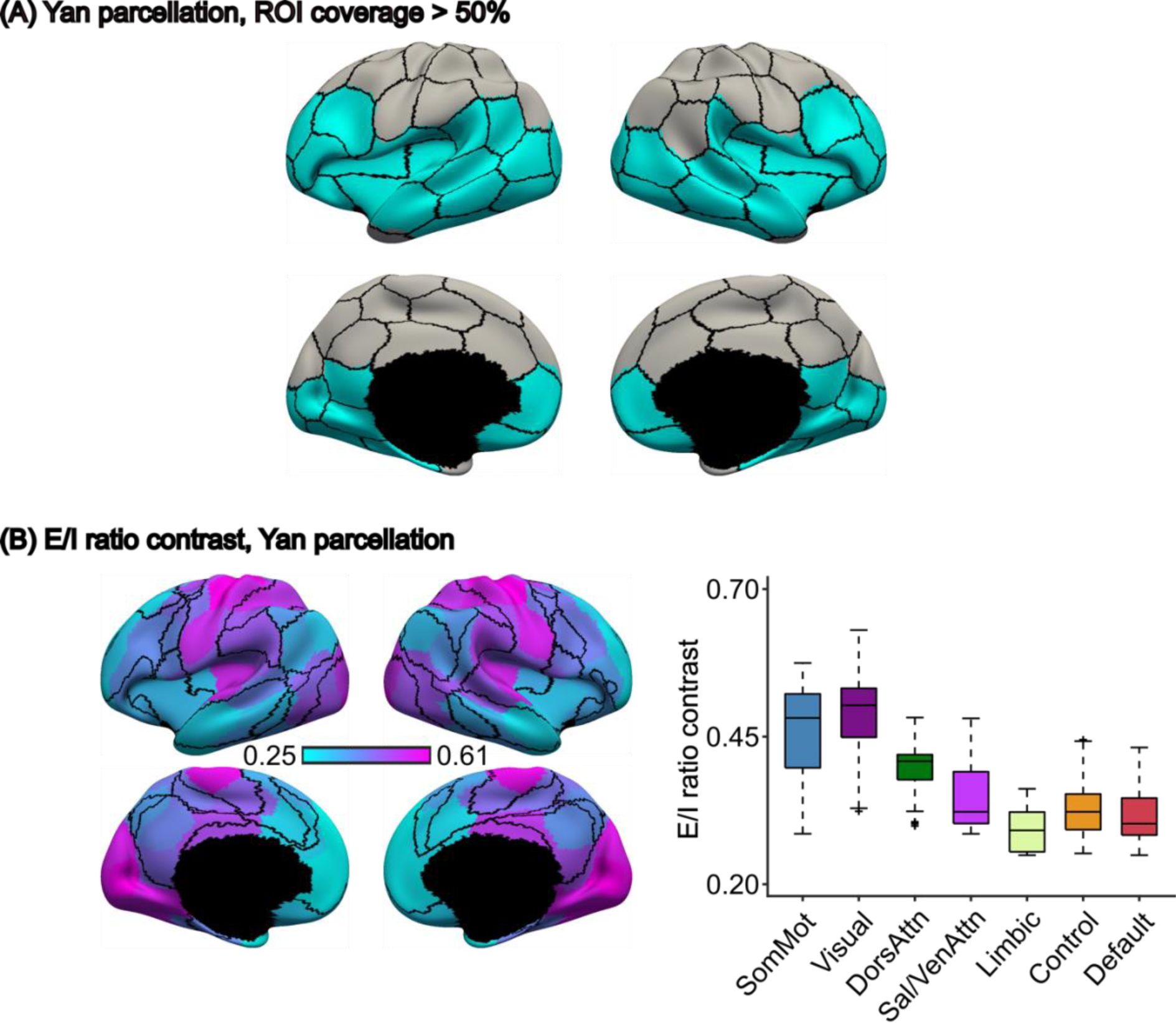
(A) Yan parcellation with 50% ROI coverage threshold, 51 out of 100 ROIs are left. (B) As a control analysis of the alprazolam E/I ratio contrast result, we changed the parcellation scheme to a higher-resolution 100-ROI parcellation. Yan parcellation has ROIs that are symmetric for the left and right hemispheres. We repeated the same analysis, and the model parameters were retrained using the same optimization approach. The boxes show the inter-quartile range (IQR) and the median. Whiskers indicate 1.5 IQR. Black crosses represent outliers.

**Figure S11.**
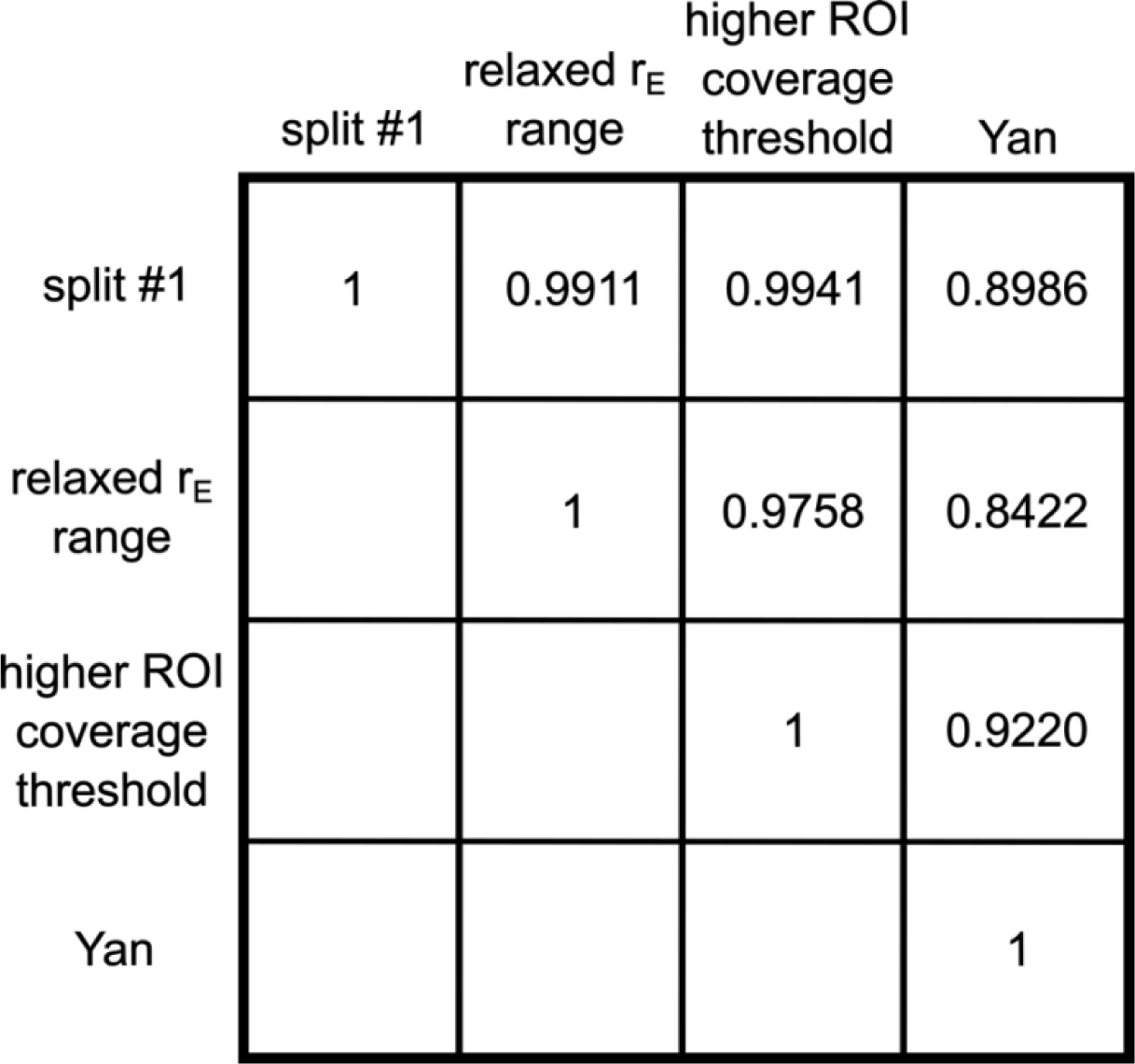
Pairwise correlation between the regional E/I ratio contrast across different control analyses based on split #1. Split #1 corresponds to the results shown in Figure 3 of the main text. The spatial distribution of region E/I ratio contrast are highly similar across different control analyses (*r* = 0.9373 ± 0.0606, mean ± std). Only the upper triangle of the matrix is shown.

**Figure S12.**
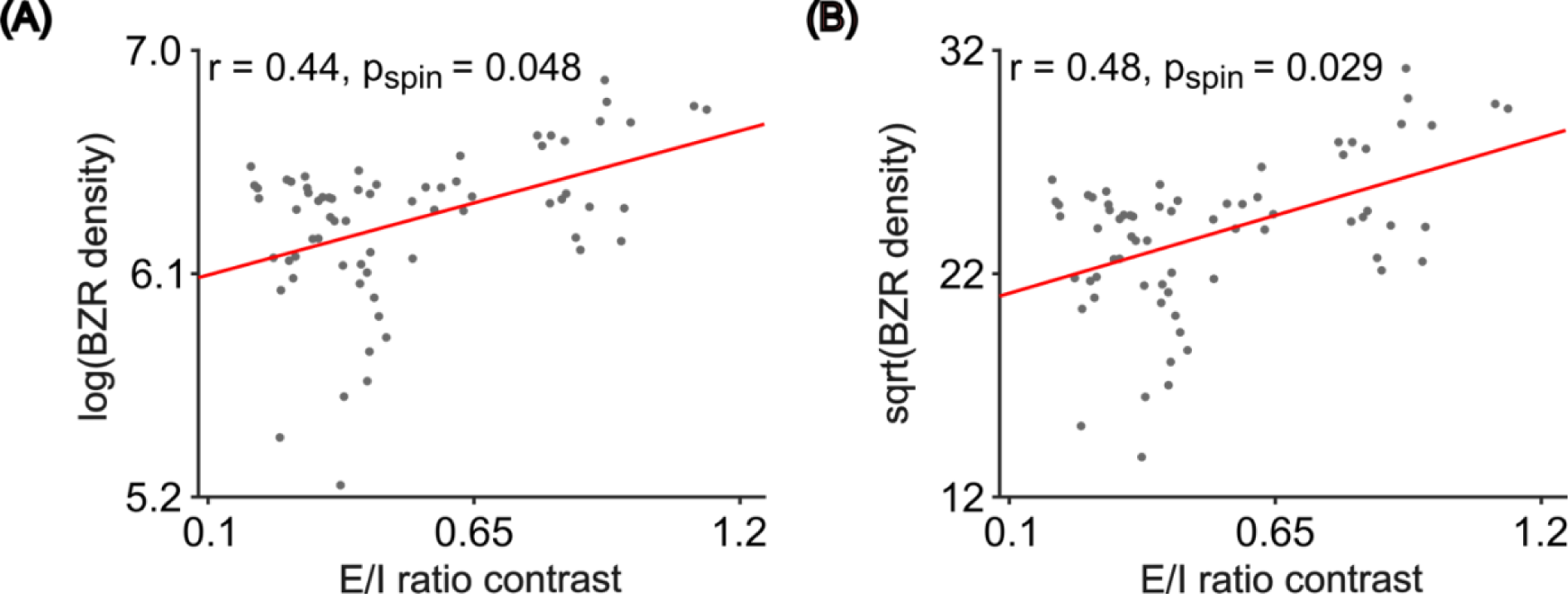
(A) Spatial correlation between regional E/I ratio contrast and log-transformed benzodiazepine receptor (BZR) density (*r* = 0.44). (B) Spatial correlation between regional E/I ratio contrast and the square root of BZR density (*r* = 0.48). Both correlations were weaker than the main results (Figure 3D), although the correlations remained statistically significant.

**Figure S13.**
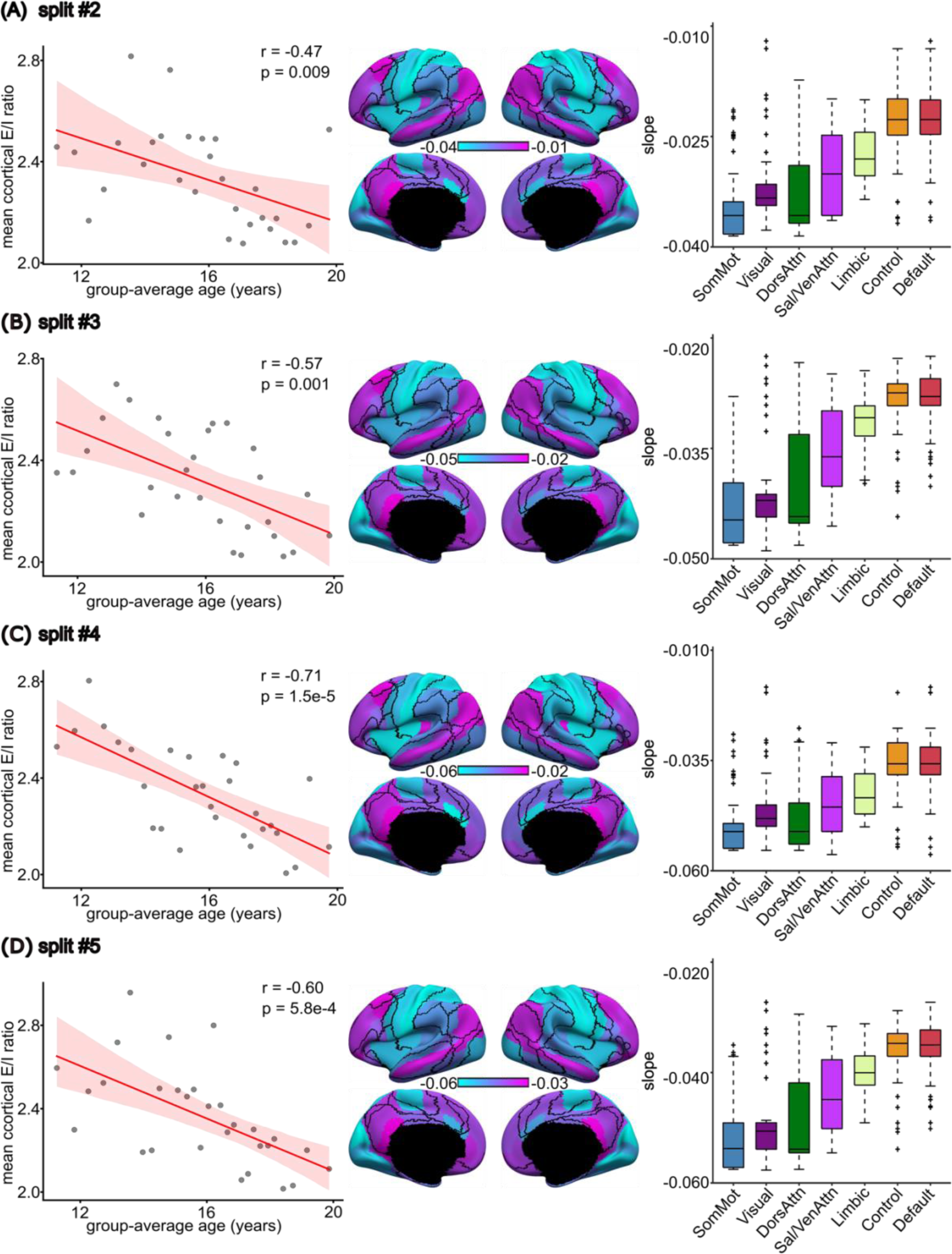
(A to D) PNC developmental analysis results obtained from 4 additional training-validation participants splits. (Left) The mean cortical E/I ratio decreases with increasing age. (Middle) The spatial distribution of regional rate of E/I ratio reduction. (Right) The regional rate of E/I ratio reduction followed a hierarchical sensorimotor-association (S-A) axis. To generate a training-validation split, 885 PNC participants were sorted according to age in an ascending order and divided into 29 age groups. Within each age groups, participants were randomly assigned to training and validation sets. The random participant splits and analyses were repeated 5 times (One shown in Figure 4, the other four shown here). All 5 splits exhibited the similar pattern of overall E/I ratio reduction and spatial distribution of the rate of reduction. We chose the split that had the highest median correlation of regional rate of E/I ratio reduction with the other 4 splits as the most representative split and showed in the main result section. The shaded area depicts 95% confidence interval of the linear relationship. The boxes show the inter-quartile range (IQR) and the median. Whiskers indicate 1.5 IQR. Black crosses represent outliers.

**Figure S14.**
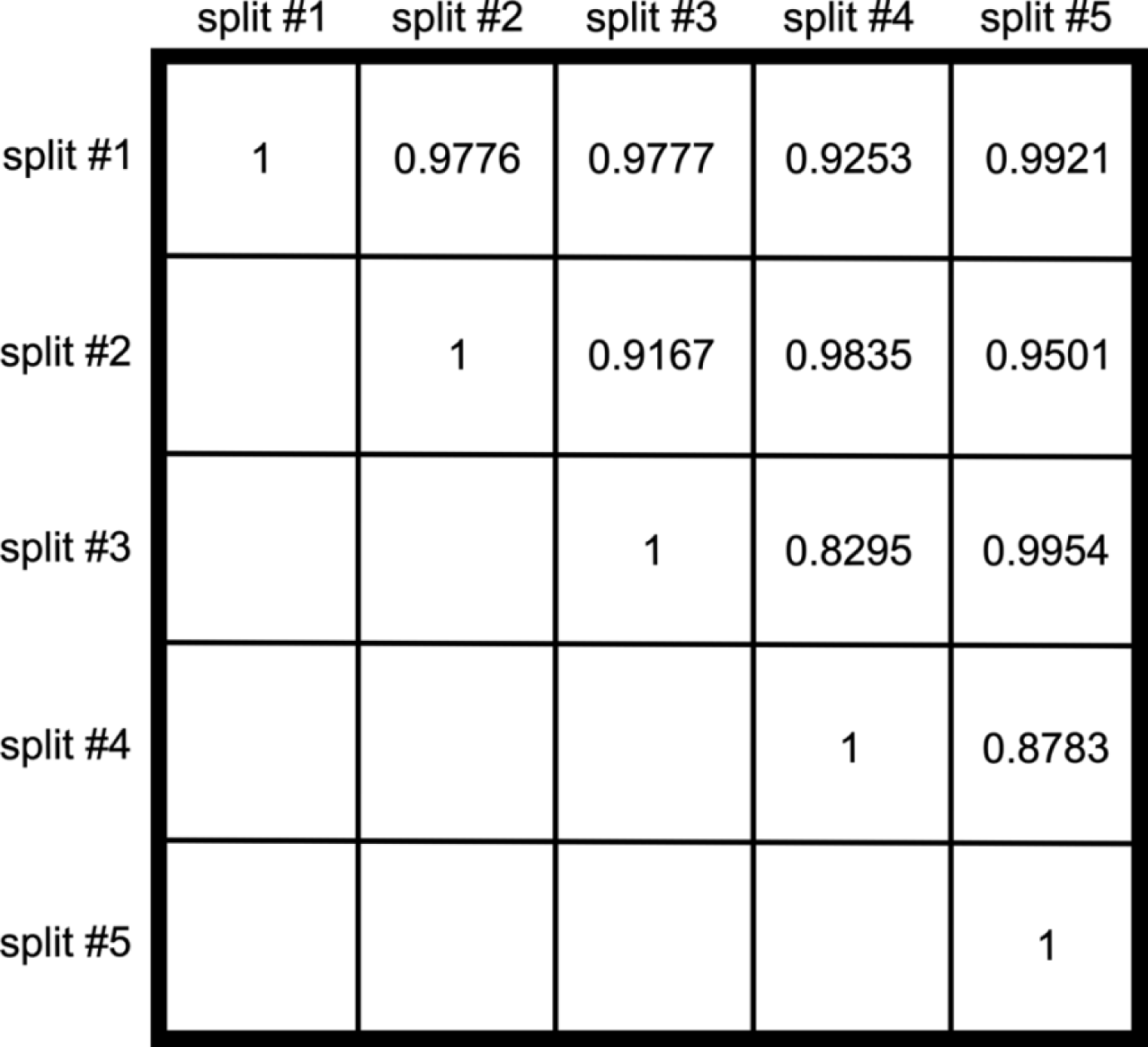
Pairwise correlation between the regional rate of E/I ratio reduction of 5 training-validation participant splits. Split #1 corresponds to the results shown in Figure 4 of the main text. The spatial distributions of the rate of E/I ratio reduction are highly similar across the 5 splits (*r* = 0.9426 ± 0.0551, mean ± std). The surface maps of different splits are shown in Figure S13. Only the upper triangle of the matrix is shown.

**Figure S15.**
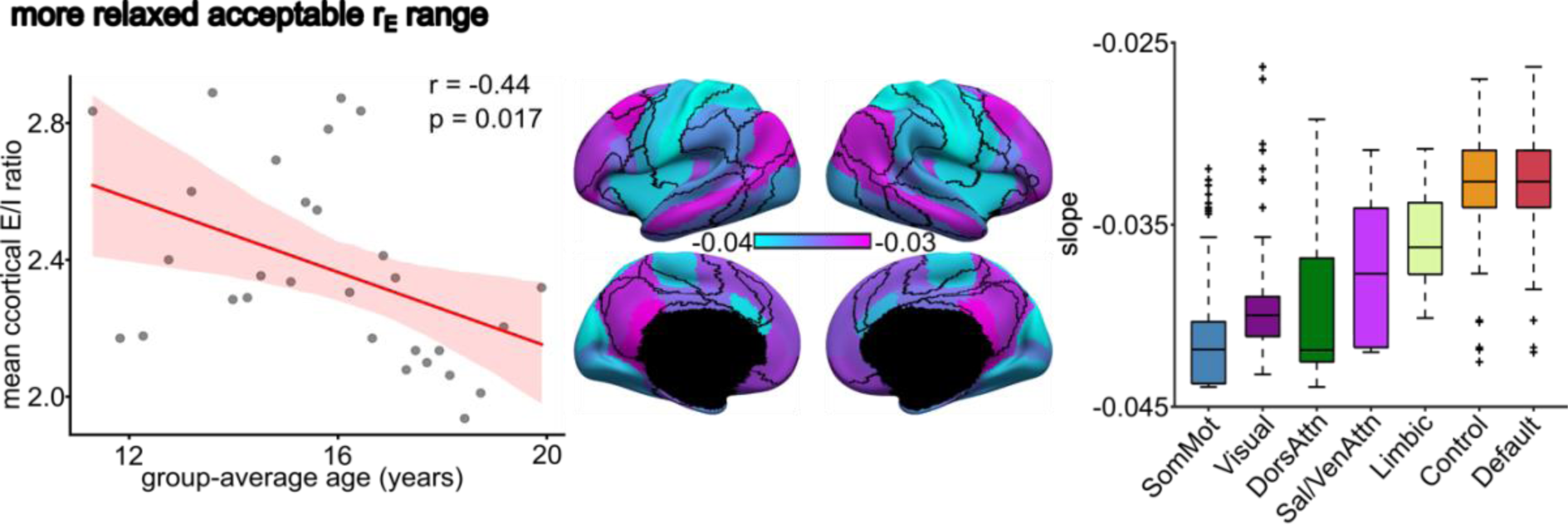
PNC developmental analysis results with relaxed excitatory firing rates thresholds. This figure is similar to Figure 4 but with the acceptable excitatory firing rates range was set to be less strict (2.5Hz to 3.5Hz). (Left) The scatter plot of overall E/I ratio reduction during development. The mean cortical E/I ratio is the average E/I ratio across all ROIs. (Middle) The spatial distribution of region E/I ratio reduction rate. (Right) Box plot of the vertex-level E/I ratio grouped by 7 resting-state networks. The boxplots comprised values obtained by “transferring” the parameter estimates from the 68 Desikan parcels to all vertices (from the underlying cortical meshes) comprising each anatomical parcel. The vertex wise parameter values were then segregated based on the seven resting-state networks. Therefore, there were 3203, 2478, 1523, 1520, 1067, 1438 and 2886 values comprising the boxplots for somatomotor, visual, dorsal attention, ventral attention, limbic, control and default networks respectively. The shaded area depicts 95% confidence interval of the linear relationship. The boxes show the inter-quartile range (IQR) and the median. Whiskers indicate 1.5 IQR. Black crosses represent outliers. The rate of E/I ratio reduction follows the sensorimotor-association (S-A) axis. E/I ratio exhibits the fastest rate of reduction in sensory regions compared to association regions.

**Figure S16.**
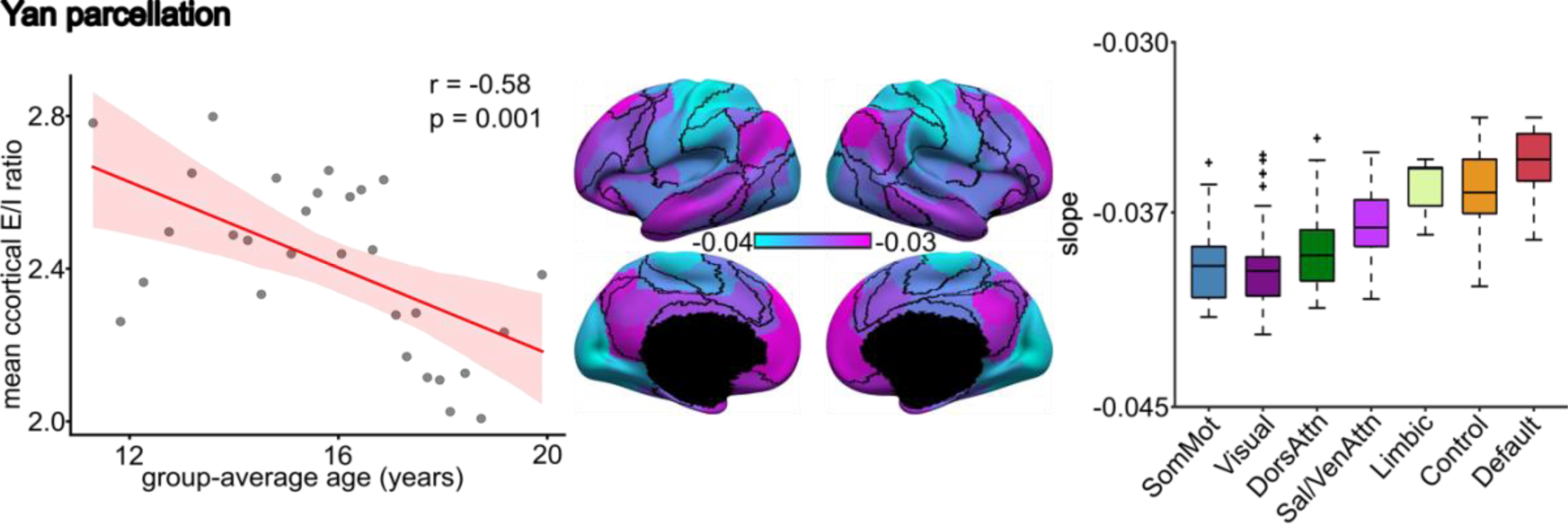
PNC developmental analysis results in Yan 100-ROI parcellation. This figure is similar to Figure 4 but utilizes the Yan 100-ROI parcellation with symmetric left and right hemisphere ROIs. (Left) The scatter plot of overall E/I ratio reduction during development. The mean cortical E/I ratio is the average E/I ratio across all ROIs. (Middle) The spatial distribution of region E/I ratio reduction rate. (Right) Box plot of the vertex-level E/I ratio grouped by 7 resting-state networks. The boxplots comprised values obtained by “transferring” the parameter estimates from the 100 Yan parcels to all vertices (from the underlying cortical meshes) comprising each anatomical parcel. The vertex wise parameter values were then segregated based on the seven resting-state networks. Therefore, there were 3203, 2478, 1523, 1520, 1067, 1438 and 2886 values comprising the boxplots for somatomotor, visual, dorsal attention, ventral attention, limbic, control and default networks respectively. The shaded area depicts 95% confidence interval of the linear relationship. The boxes show the inter-quartile range (IQR) and the median. Whiskers indicate 1.5 IQR. Black crosses represent outliers. The rate of E/I ratio reduction follows the sensorimotor-association (S-A) axis. E/I ratio exhibits the fastest rate of reduction in sensory regions compared to association regions.

**Figure S17.**
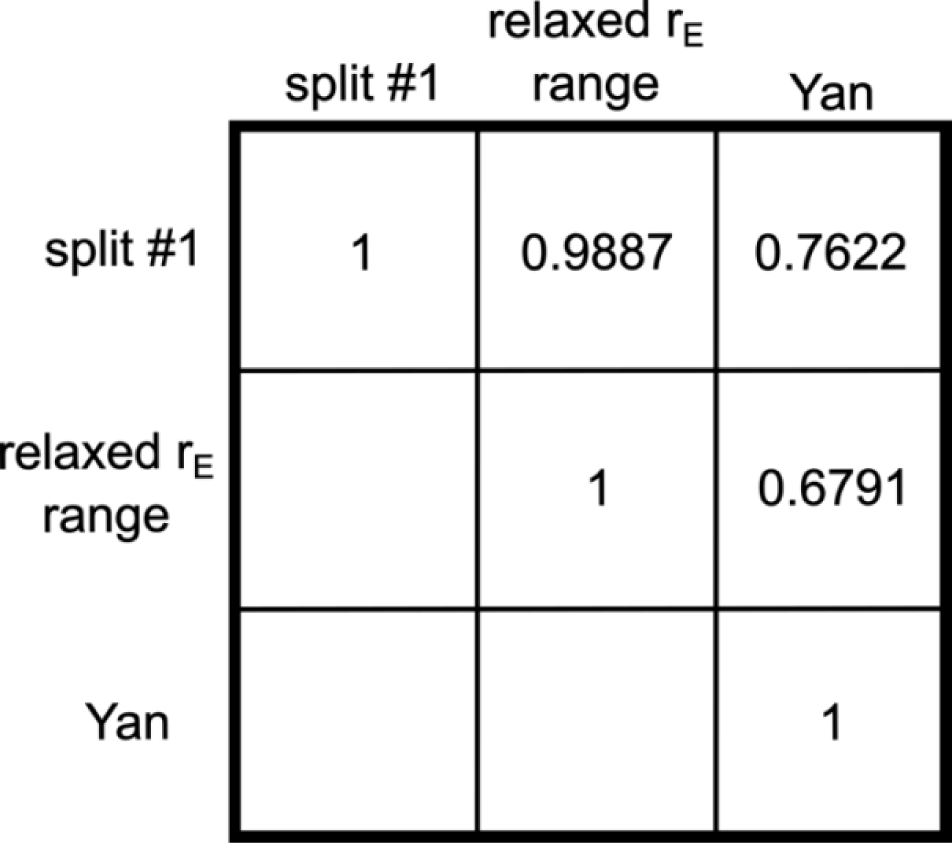
Pairwise correlation between the regional rate of E/I ratio reduction of different control analyses based on split #1. Split #1 corresponds to the results shown in Figure 4 of the main text. The spatial distributions of the rate of E/I ratio reduction are highly similar across different control analyses (*r* = 0.8100 ± 0.1603, mean ± std). The surface maps of different cases are shown in the supplementary Figure S15 and S16. Only the upper triangle of the matrix is shown.

**Figure S18.**
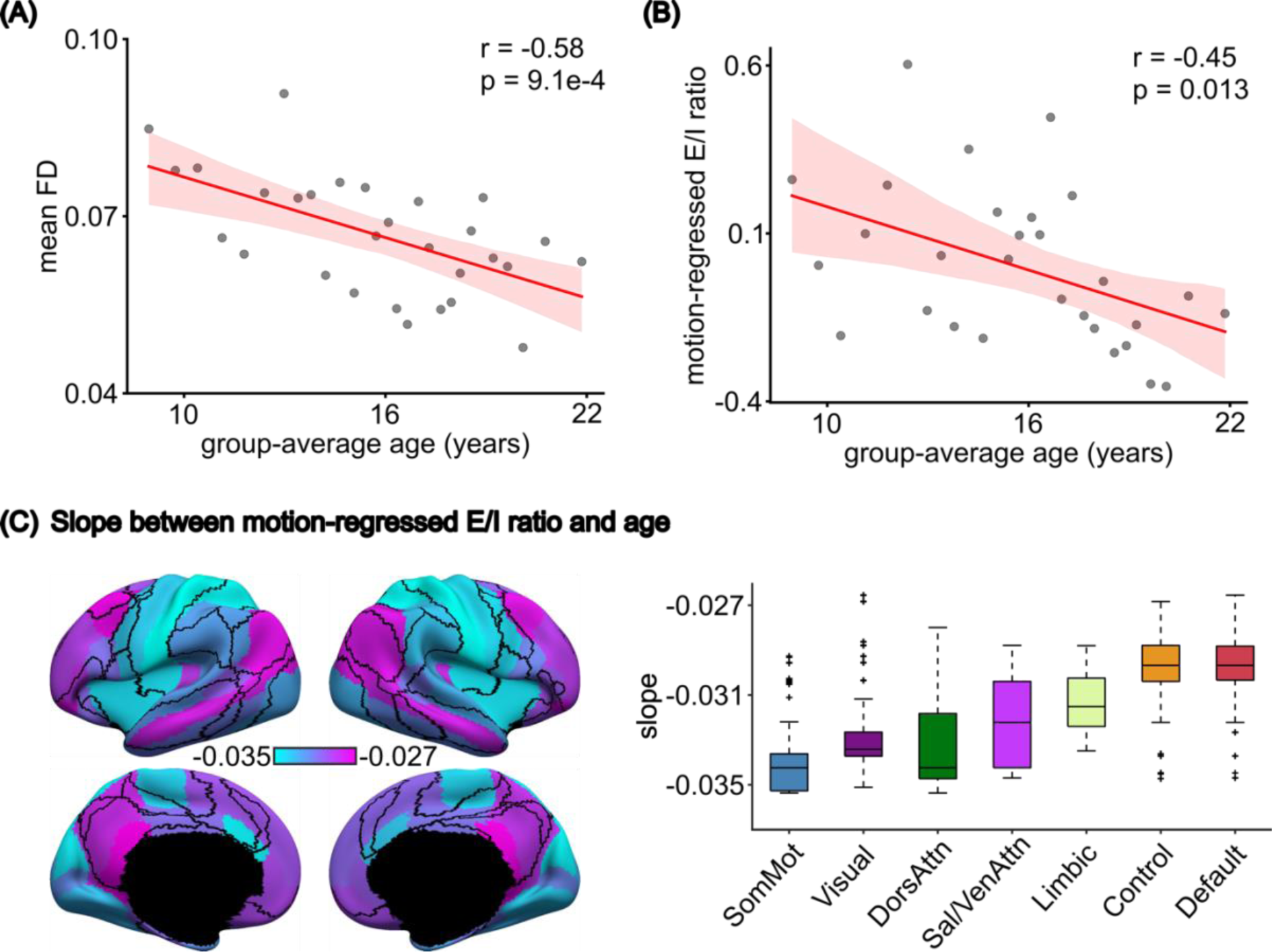
(A) Mean framewise displacement (FD) decreased with age (*r* = −0.58, *p* = 9.1e-4). (B) After regressing out mean FD from the estimated E/I ratio across all age groups, the residuals still significantly decreased with age (*r* = −0.45, *p* = 0.013). (C) Spatial distribution of linear regression slope between FD-regressed E/I ratio and age. All slopes were negative and significant (FDR *q* < 0.05). (D) The slopes exhibited a spatial gradient with sensory-motor networks showing the fastest reduction and association networks showing slower reduction. The boxes show the inter-quartile range (IQR) and the median. Whiskers indicate 1.5 IQR. Black crosses represent outliers.

**Figure S19.**
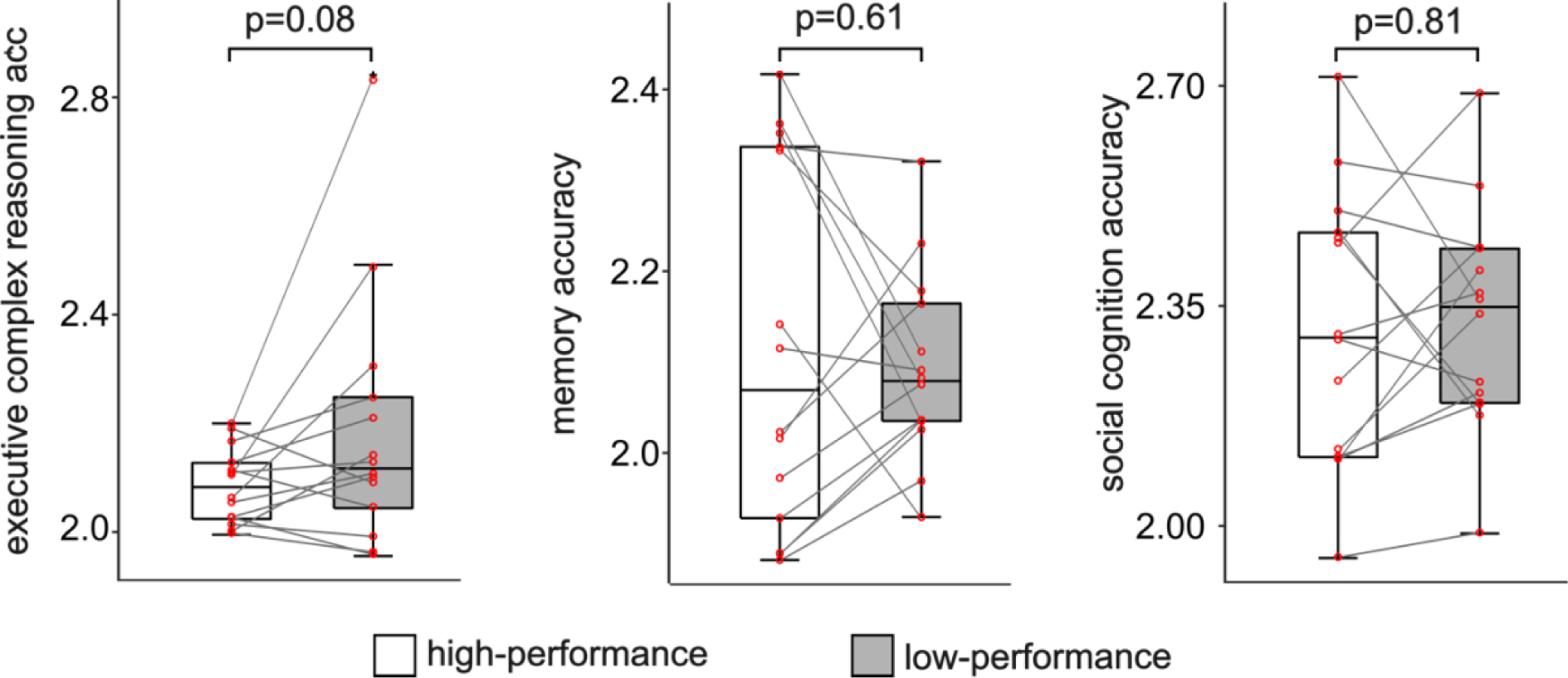
Box plots of E/I ratio estimated from the three domain-specific accuracy scores from Penn Computerized Neurocognitive Battery (CNB). For each 1-year interval, participants whose age are within this interval are extracted to form one age group. For each age group and cognitive score, participants with cognitive scores above the median are assigned to a high-performance group, the rest are assigned to a low-performance group. Both high- and low-performance groups are further divided into subgroups. Participants within each subgroup are randomly assigned to training and validation sets. Each box plot shows the E/I ratio of high- and low-performance group associated with each cognitive score. We observed no significant difference between E/I ratio of high- and low-performance groups for any of the three cognitive scores. The boxes show the inter-quartile range (IQR) and the median. Whiskers indicate 1.5 IQR.

**Figure S20.**
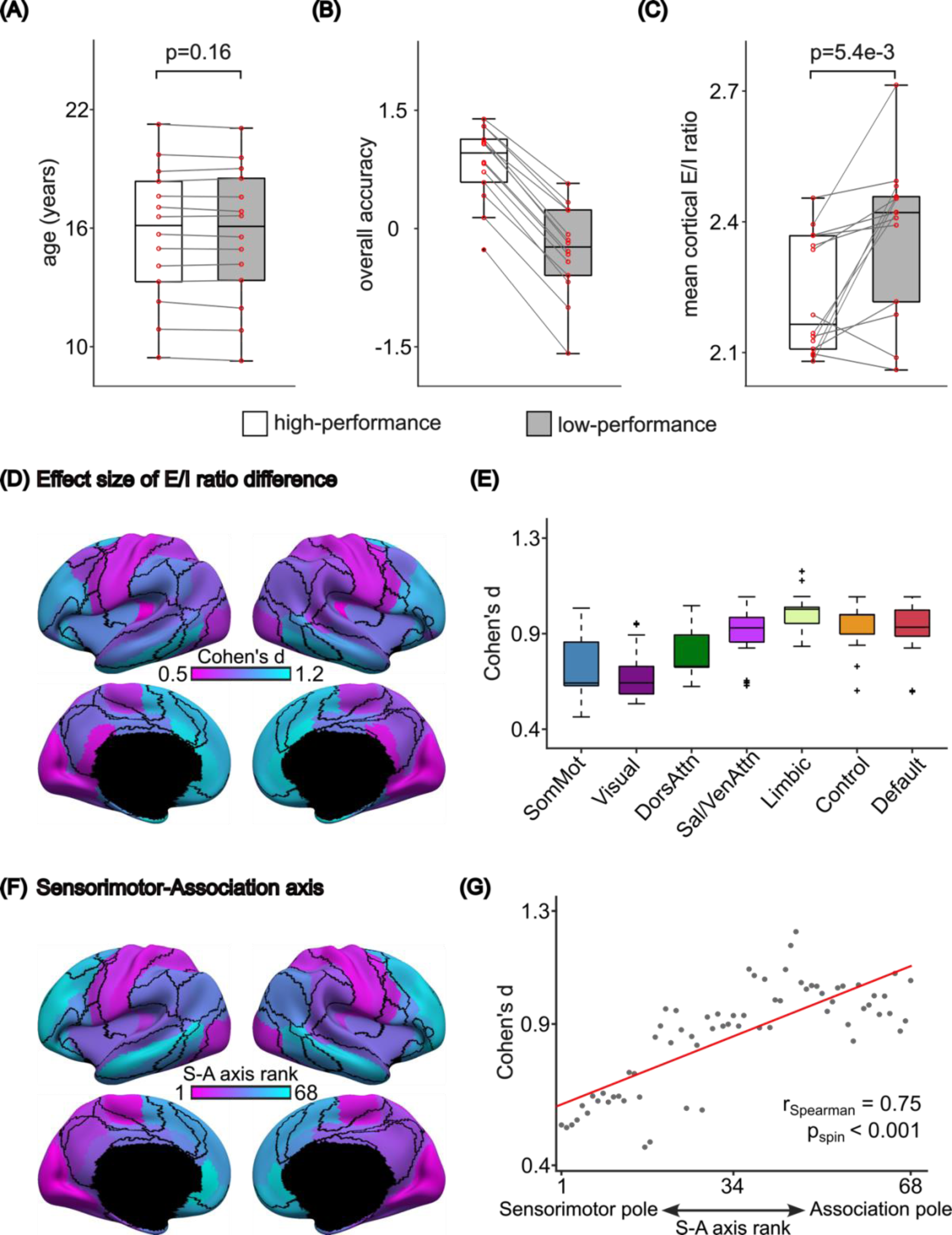
PNC cognition analysis results obtained from 4 additional training-validation participants splits (Figure. S20 to S23). To generate a training-validation split, 885 PNC participants were sorted according to age in an ascending order. For each 1-year interval, participants whose age are within this interval are extracted to form one age group. For each age group, participants with cognitive scores above the median are assigned to a high-performance group, the rest are assigned to a low-performance group. Both high- and low-performance groups are further divided into 14 subgroups. Participants within each subgroup are randomly assigned to training and validation sets. The random participant splits and analyses were repeated 5 times. (A) Boxplots of age, (B) ‘Overall accuracy’, and (C) Mean cortical E/I ratio of high- and low-performance groups. (D) Spatial distribution of effect size of regional E/I ratio difference between high-performance and low-performance groups. (E) On a network level, the effect sizes of E/I ratio differences follow a hierarchical structure. We chose the split that had the highest median correlation of regional Cohen’s d values with the other 4 splits as the most representative split and showed in the main result section. The results of the 4 splits are consistent with our main results. The boxes show the inter-quartile range (IQR) and the median. Whiskers indicate 1.5 IQR. (F) ROI rankings along the sensorimotor-association (S-A) axis. Lower ranks were assigned to ROIs that were more towards the sensorimotor pole; higher ranks were assigned to ROIs that were more towards the association pole. (G) Agreement between the effect size of E/I ratio difference and S-A axis rank.

**Figure S21.**
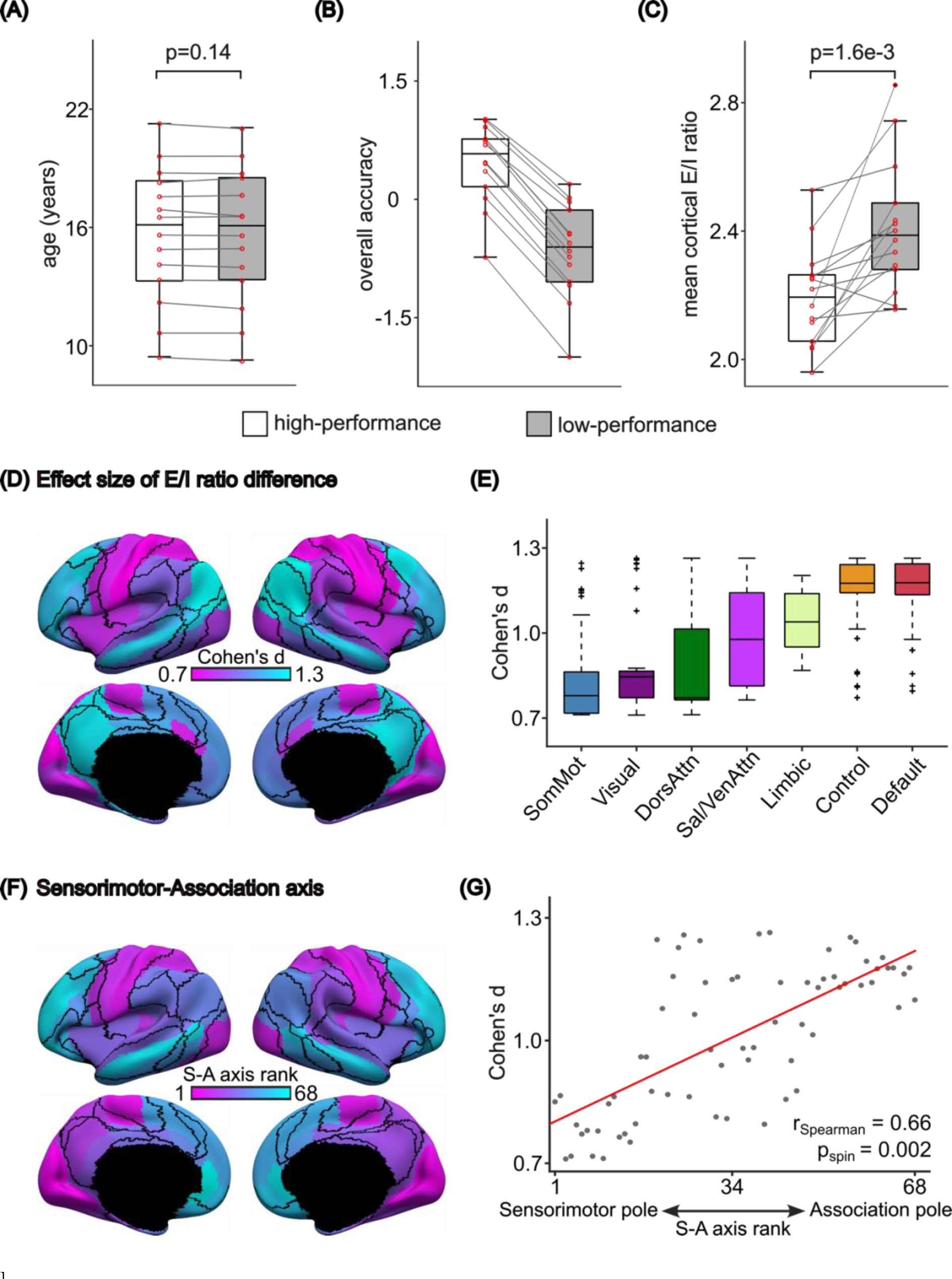
The 3rd training-validation participant split for PNC cognition analysis.

**Figure S22.**
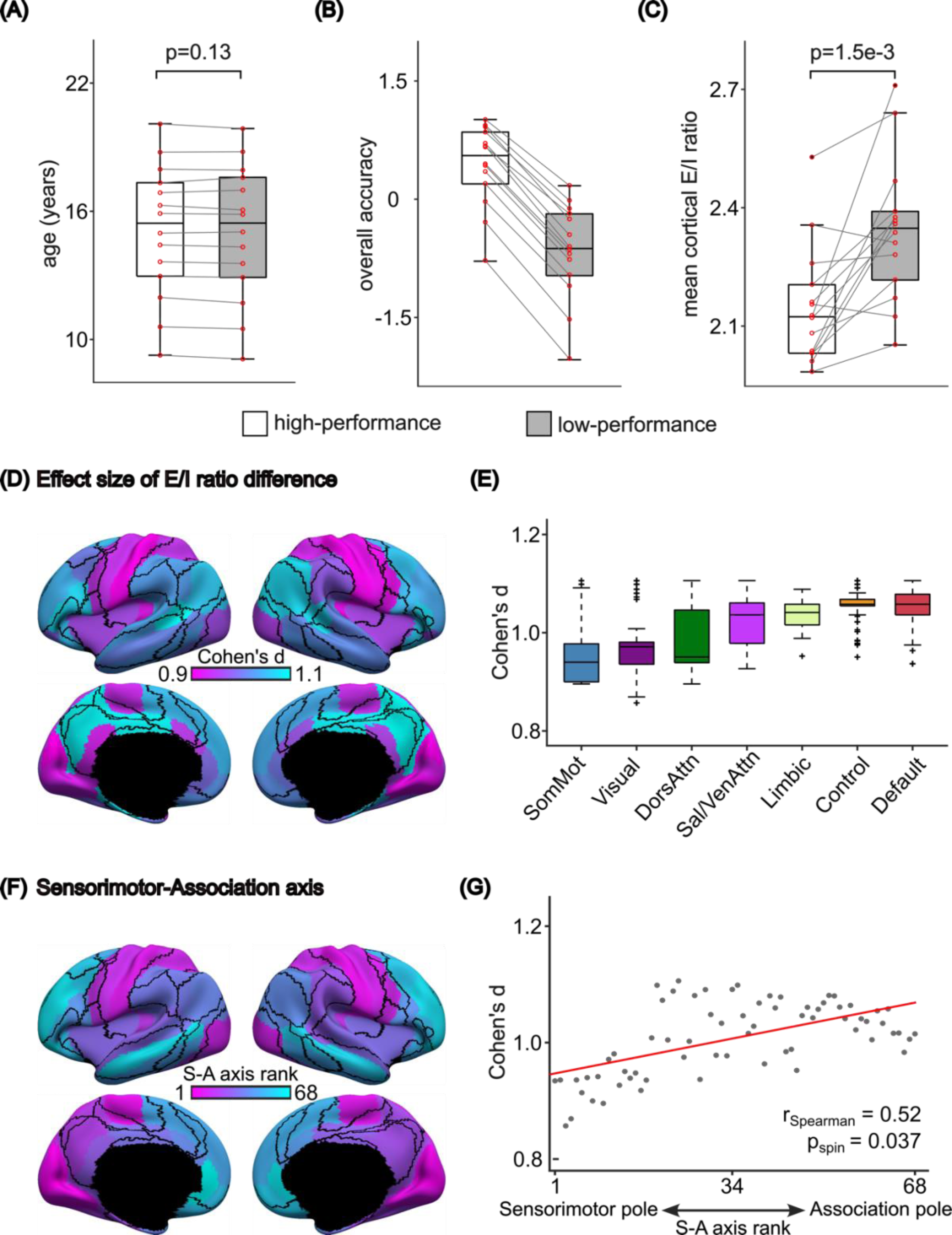
The 4th training-validation participant split for PNC cognition analysis.

**Figure S23.**
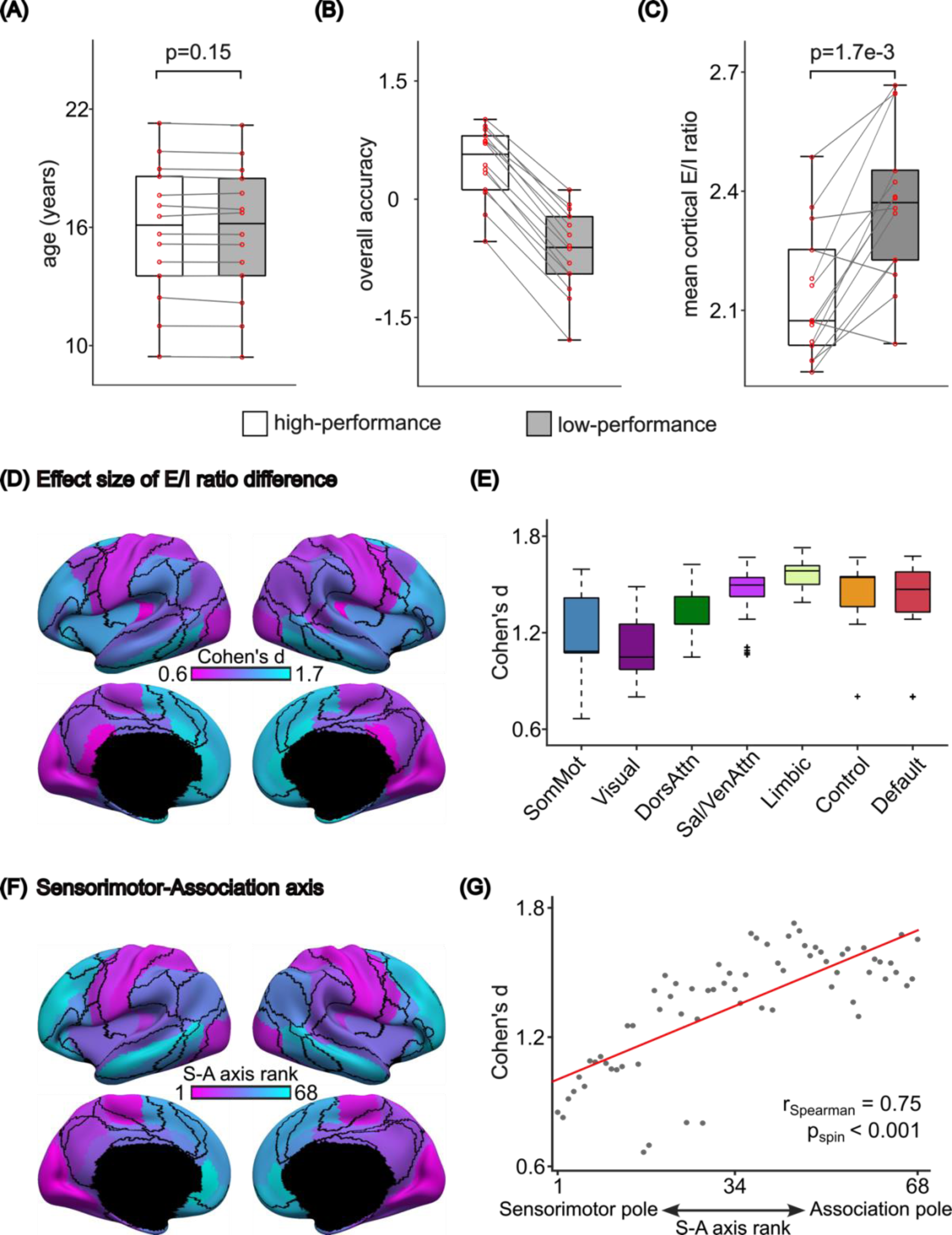
The 5th training-validation participant split for PNC cognition analysis.

**Figure S24.**
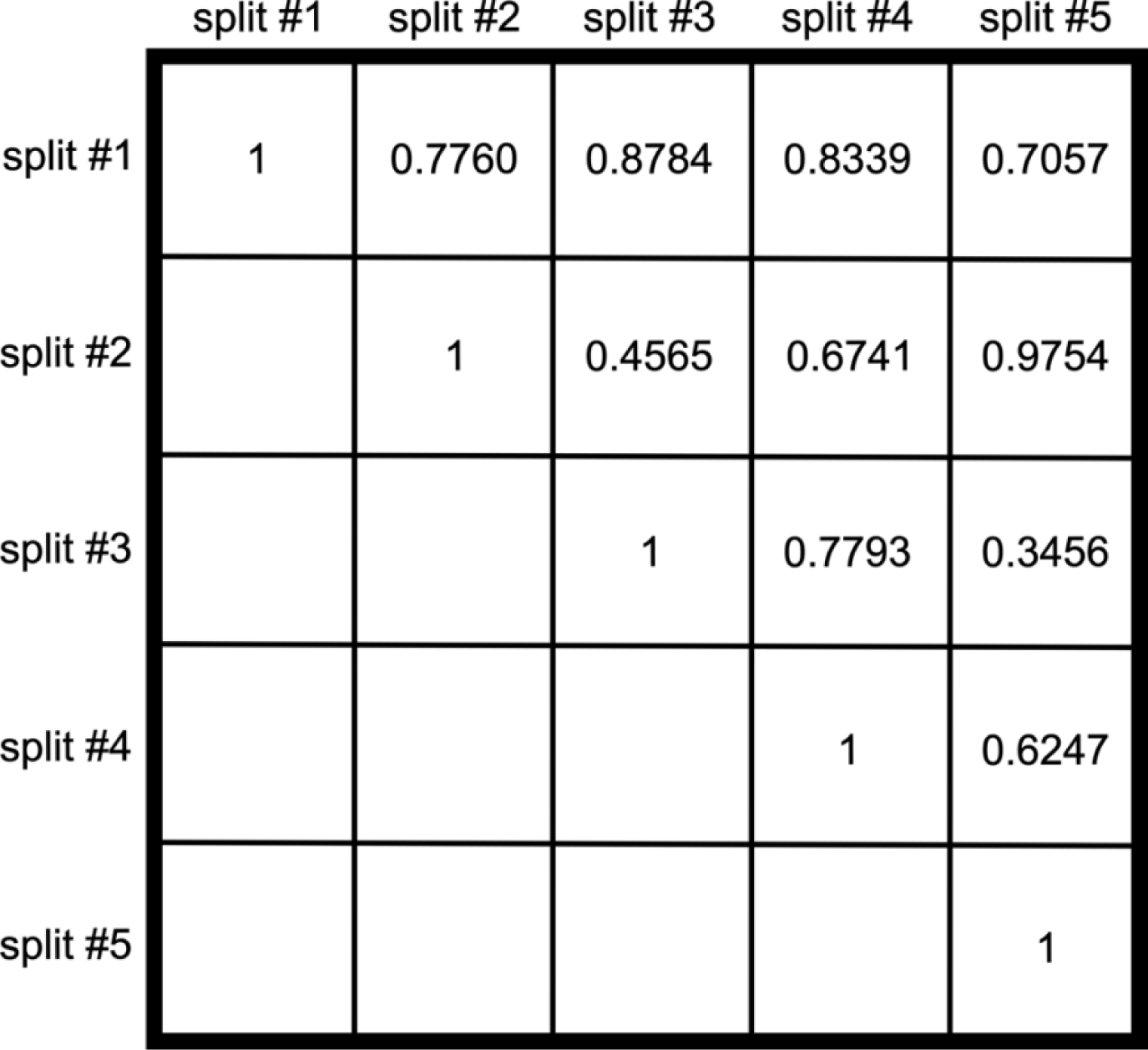
Pairwise correlation of regional effect sizes (i.e., Cohen’s d) between E/I ratios of high- and low-performance groups across 5 participant splits (split #1-5). For each split, high- and low-performance groups were divided into 14 subgroups with training and validation sets. The spatial distributions of the rate of E/I ratio reduction are highly similar across the 5 splits (*r* = 0.7050 ± 0.1911, mean ± std). Split #1 corresponds to the results shown in Figure 5 of the main text. The surface maps of different splits are shown in the main result (Figure 5D) and supplementary information (Figure. S20 to S23). Only the upper triangle of the matrix is shown.

**Figure S25.**
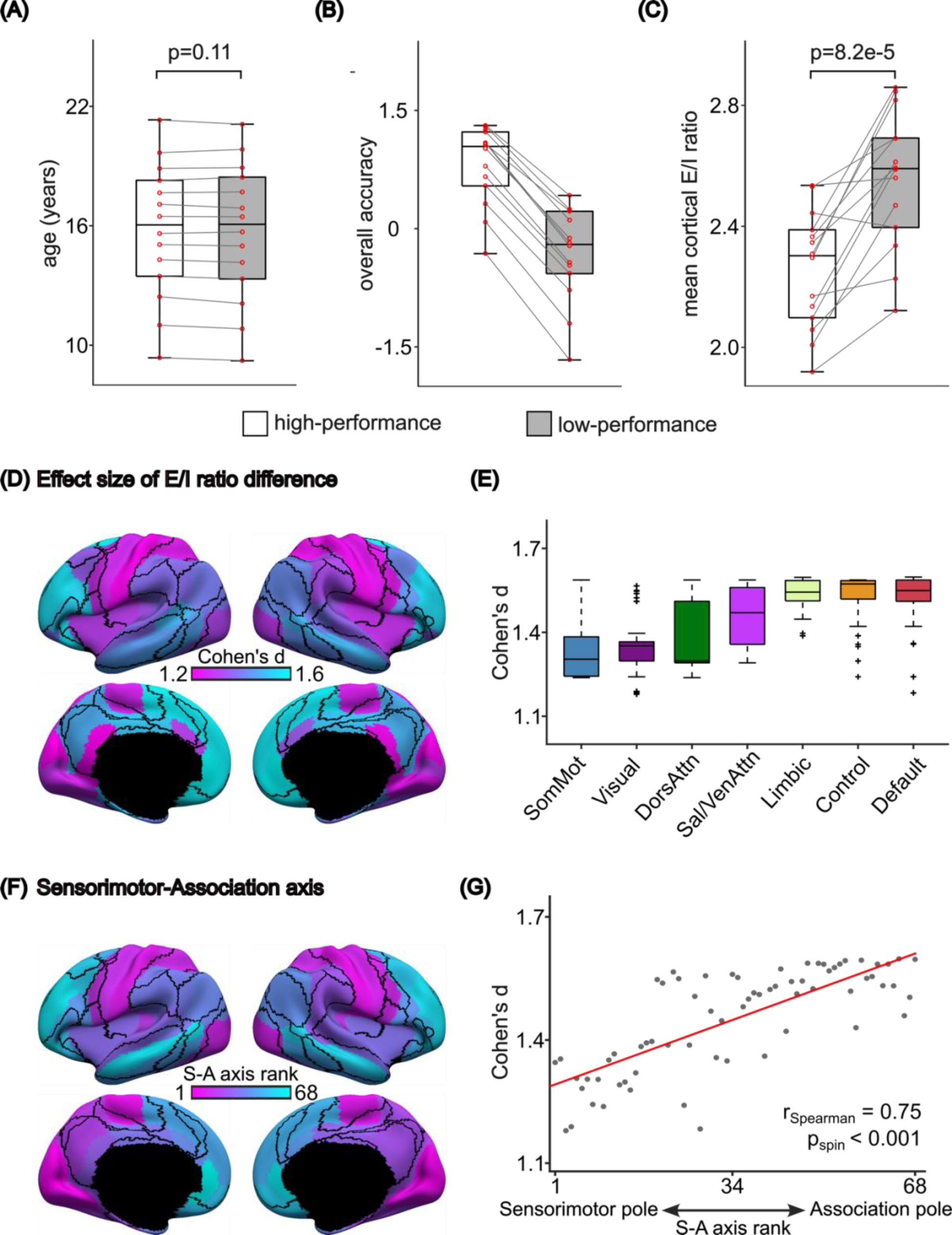
PNC cognition analysis results with relaxed excitatory firing rates thresholds. This figure is similar to Figure 5 but with the acceptable excitatory firing rates range was set to be less strict (2.5Hz to 3.5Hz). (A) Boxplots of age, (B) ‘Overall accuracy’, and (C) Mean cortical E/I ratio of high- and low-performance groups. (D) Spatial distribution of effect size of regional E/I ratio difference between high-performance and low-performance groups. (E) Box plot of Cohen’s d of vertex-level E/I ratio differences grouped by 7 resting-state networks. The boxplots comprised values obtained by “transferring” the parameter estimates from the 68 Desikan parcels to all vertices (from the underlying cortical meshes) comprising each anatomical parcel. The vertex wise parameter values were then segregated based on the seven resting-state networks. Therefore, there were 3203, 2478, 1523, 1520, 1067, 1438 and 2886 values comprising the boxplots for somatomotor, visual, dorsal attention, ventral attention, limbic, control and default networks respectively. The boxes show the inter-quartile range (IQR) and the median. Whiskers indicate 1.5 IQR. Black crosses represent outliers. The difference of E/I ratio between high- and low-performance groups follows a hierarchical structure. Cohen’s d of E/I ratio differences in cognition is larger in association regions compared to sensory regions. (F) ROI rankings along the sensorimotor-association (S-A) axis. Lower ranks were assigned to ROIs that were more towards the sensorimotor pole; higher ranks were assigned to ROIs that were more towards the association pole. (G) Agreement between the effect size of E/I ratio difference and S-A axis rank.

**Figure S26.**
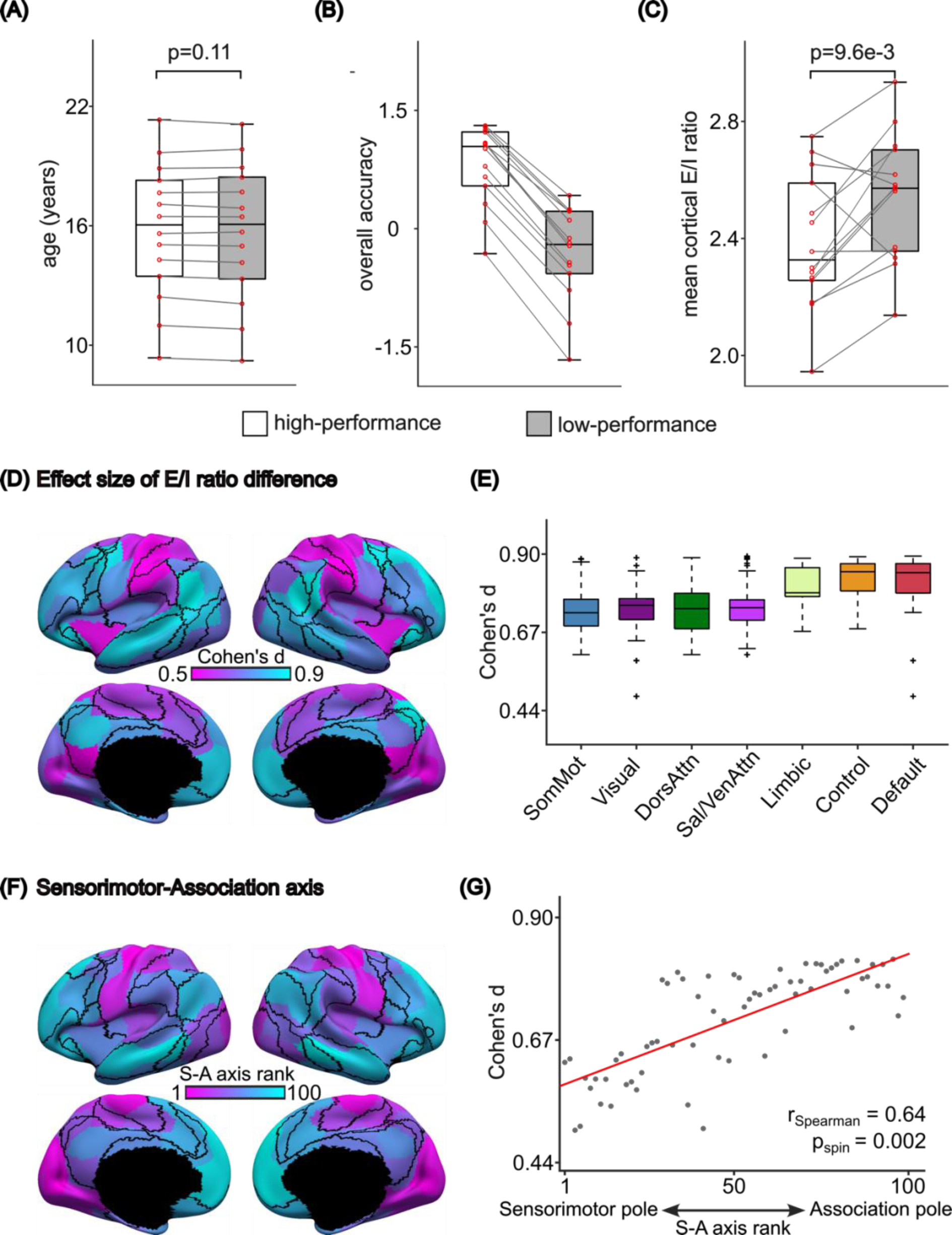
PNC cognition analysis results in Yan 100-ROI parcellation. This figure is similar to Figure 5 but utilizes the Yan 100-ROI parcellation with symmetric left and right hemisphere ROIs. (A) Boxplots of age, (B) ‘Overall accuracy’, and (C) Mean cortical E/I ratio of high- and low-performance groups. (D) Spatial distribution of effect size of regional E/I ratio difference between high-performance and low-performance groups. (E) Box plot of Cohen’s d of vertex-level E/I ratio differences grouped by 7 resting-state networks. The boxplots comprised values obtained by “transferring” the parameter estimates from the 100 Yan parcels to all vertices (from the underlying cortical meshes) comprising each anatomical parcel. The vertex wise parameter values were then segregated based on the seven resting-state networks. Thepre, there were 3203, 2478, 1523, 1520, 1067, 1438 and 2886 values comprising the boxplots for somatomotor, visual, dorsal attention, ventral attention, limbic, control and default networks respectively. The boxes show the inter-quartile range (IQR) and the median. Whiskers indicate 1.5 IQR. Black crosses represent outliers. The difference of E/I ratio between high- and low-performance groups follows the sensory-to-association (SA) axis. Cohen’s d of E/I ratio differences in cognition is larger in association regions compared to sensory regions. (F) ROI rankings along the sensorimotor-association (S-A) axis. Lower ranks were assigned to ROIs that were more towards the sensorimotor pole; higher ranks were assigned to ROIs that were more towards the association pole. (G) Agreement between the effect size of E/I ratio difference and S-A axis rank.

**Figure S27.**
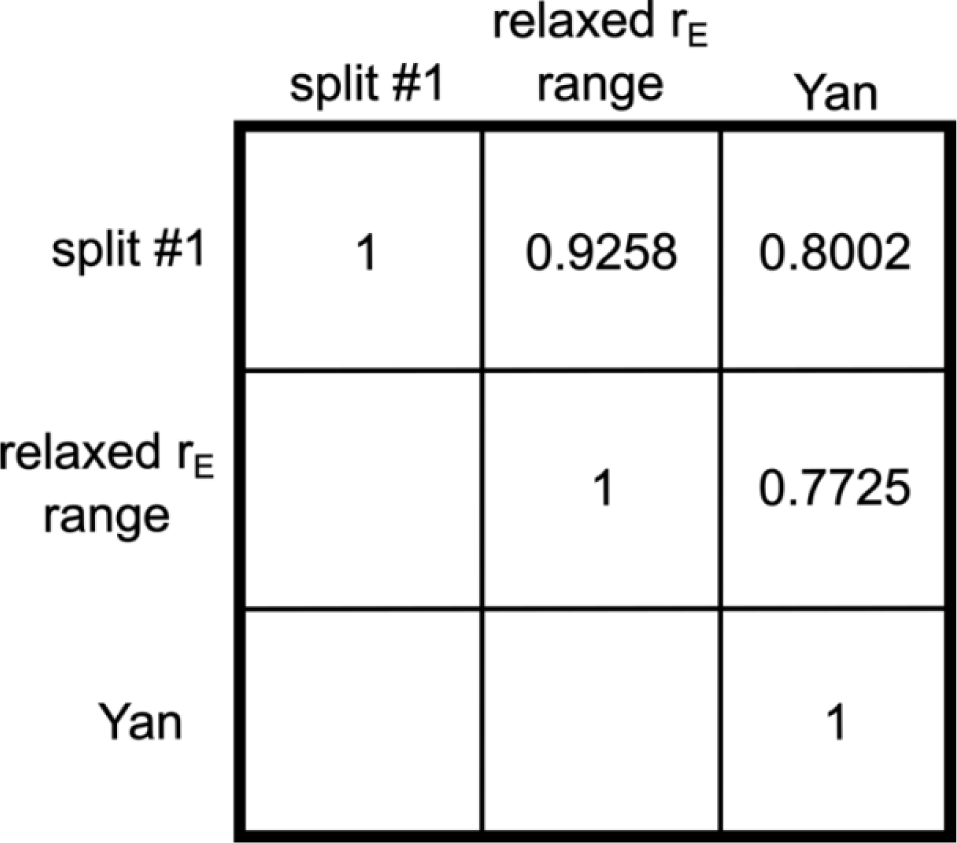
Pairwise correlation of regional effect sizes (i.e., Cohen’s d) between E/I ratios of high- and low-performance groups across different control analyses based on split #1. Split #1 corresponds to the results shown in Figure 5 of the main text. The spatial distributions of the rate of E/I ratio reduction are highly similar across different control analyses (*r* = 0.8328 ± 0.0817, mean ± std). Only the upper triangle of the matrix is shown.

